# Stabilization of Brain Network Dynamics during Childhood and Adolescence is Associated with Gene Expressions

**DOI:** 10.1101/2021.03.03.433828

**Authors:** Tianyuan Lei, Xuhong Liao, Xiaodan Chen, Tengda Zhao, Yuehua Xu, Mingrui Xia, Jiaying Zhang, Xiaochen Sun, Yongbin Wei, Weiwei Men, Yanpei Wang, Mingming Hu, Gai Zhao, Bin Du, Qian Wu, Shuping Tan, Jiahong Gao, Shaozheng Qin, Sha Tao, Qi Dong, Yong He

## Abstract

Functional brain networks require dynamic reconfiguration to support flexible cognitive function. However, the developmental principles shaping brain network dynamics remain poorly understood. Here, we report the longitudinal development of large-scale brain network dynamics during childhood and adolescence, and its connection with gene expression profiles. Using a multilayer network model, we show the temporally varying modular architecture of child brain networks, with higher network switching primarily in the association cortex and lower switching in the primary regions. This topographical profile exhibits progressive maturation, which manifests as reduced modular dynamics, particularly in the transmodal (e.g., default-mode and frontoparietal) and sensorimotor regions. These developmental refinements mediate age-related enhancements of global network segregation and are linked with the expression profiles of genes associated with the enrichment of ion transport and nucleobase-containing compound transport. These results highlight a progressive stabilization of brain dynamics, which expand our understanding of the neural mechanisms that underlie cognitive development.

## Introduction

The human brain is an efficient and dynamic information processing system with a complex spatiotemporal organization. Network science approaches have revealed that the human brain functional network contains a non-trivial modular structure, in which functional specialization and integration are well balanced at low wiring cost (He et al., 2009; Liao et al., 2017; Meunier et al., 2010; Sporns & Betzel, 2016). This modular structure facilitates a fast response to domain-specific stimuli (Bertolero et al., 2015; Meunier et al., 2010; Sporns & Betzel, 2016) and also enables efficient global brain communication (Bullmore & Sporns, 2012). Recent studies suggest that the modular organization of the brain is not static but rather shows temporally varying patterns over short time scales (e.g., seconds), with higher network switching primarily in the association (e.g., frontoparietal) regions and lower variability in the primary regions (Chen et al., 2016; Liao et al., 2017; Liu et al., 2020; Pedersen et al., 2018). These network dynamics contribute significantly to flexible cognitive function (Chen et al., 2016; Liao et al., 2017; Pedersen et al., 2018; Yin et al., 2020) and are distinctive to each individual (Liao et al., 2017). During short-term training, network segregation in the brain is increased through dynamic reconfigurations, and these increases correspond to improved task automation (Bassett et al., 2015; Finc et al., 2020). These findings suggest a link between adaptive network dynamics and skill acquisition which we believe may be significant not only during short term learning but also in long term development. However, how network dynamics changes during childhood and adolescence, a crucial stage for cognitive and behavioral development, remains largely unknown. The goal of the current study was to gain insight into the principles shaping the maturation of network dynamics in the human brain, and its connection with cognitive development and gene expression profiles.

Childhood and adolescence are critical developmental phases for the consolidation and refinement of individual motor, cognitive, social, and emotional capabilities (National research council (US), 1984; Paus et al., 2008). These physical, psychological and cognitive developments occur in parallel with the substantial development of brain architecture (Cao et al., 2016; Vértes & Bullmore, 2015). During this period, brain microstructure is fine-tuned through processes including regressive synaptic pruning and progressive myelination (Tau & Peterson, 2010; Vértes & Bullmore, 2015). At the macroscopic level, the modular structure of the functional brain networks undergoes remarkable reconfiguration with age (Cao et al., 2016; Grayson & Fair, 2017; Vértes & Bullmore, 2015), shifting from anatomical proximity during childhood to a spatially distributed layout at adulthood (Fair et al., 2009). Within modules, a decrease in the number of short-range connections with age is observed in some modules as a result of synaptic pruning (Fair et al., 2007; Supekar et al., 2009), while in others, the number of long-range connections increases over time, for example the increase in anterior-posterior connections observed in the default-mode network (Fair et al., 2008; Fan et al., 2020; Sato et al., 2014). Between modules, integration between the default-mode network and other brain systems exhibits an age-related increase, while integration between the higher-order cognitive network and the subcortical network with other brain systems show an age-related decrease (Gu et al., 2015). These system-specific changes in intra- and inter-module connections are indicative of the growing functional differentiation of brain modules with development. Notably however, previous studies on brain network modularity were mainly undertaken on the development of static (i.e., time-invariant) modular architecture, and largely ignored the temporal dynamics of brain modularity. Yet, as recent work has pointed out, cognitive growth is largely dependent on age-related adjustments in the brain’s temporal dynamics (Hutchison & Morton, 2016). To date, how modular dynamics in the brain network develops towards maturation over childhood to adolescence has yet to be established.

Structural and functional development of the brain is shaped by genetic factors (Douet et al., 2014; Johnson et al., 2009; Zhong et al., 2018). For instance, animal model studies have revealed that myelination in the central neural system is closely governed by a gene named the myelin gene regulatory factor, which is specifically expressed in oligodendrocytes (Emery et al., 2009), and synaptic pruning is mediated by astrocytes through the *Megf10* (i.e., multiple epidermal growth factor-like domains protein 10) and *Mertk* (i.e., Mer tyrosine kinase) phagocytic pathways (Chung et al., 2013). Recent advances in connectome-transcriptome association analysis now enable us to explore the transcriptional signatures underlying the spatial organization of the human brain network *in vivo* (Fornito et al., 2019). In adults, the spatial layout of functional modules is shaped by genes associated with the enrichment of ion channels (Richiardi et al., 2015). Inter-module hubs have also been shown to be metabolically expensive, with an overrepresentation of genes for oxidative metabolism and mitochondria (Vértes et al., 2016). A very recent study reported that the spatial layout of brain module dynamics in adults is associated with genes involved in potassium ion transport, establishing a link between large-scale connectivity dynamics and transcriptional profiles (Liu et al., 2020). Nevertheless, little is known on how gene expression is linked to developmental brain dynamics in children.

To address these issues, we investigated the developmental changes in brain network dynamics between childhood to adolescence, and the transcriptional profiles of genes related to this process. Brain network analyses were undertaken using a large longitudinal resting-state functional magnetic resonance imaging (rsfMRI) dataset comprised of scans from 305 healthy children (age 6-14 years, 491 scans in total) (Fan et al., 2020), and genetic analysis was conducted using postmortem gene expression data from the Allen Human Brain Atlas (Hawrylycz et al., 2012). Specifically, for all rsfMRI scans of each participant, we applied a multilayer network model (Mucha et al., 2010) to identify the time-resolved modular architecture in the child brain and further quantified the temporal switching of regional module affiliations. We aimed to investigate i) developmental patterns in brain network dynamics during childhood and adolescence at the whole-brain, system and nodal levels, and their potential association with cognitive development; ii) whether these developmental patterns contribute to age-related changes in the information transmission capability of brain networks; and iii) the association between developmental changes in brain network dynamics and gene transcriptional profiles.

## Results

### Spatial Patterns of Brain Network Dynamics in Children

We conducted a longitudinal rsfMRI study on a cohort of 305 typically developing children (age 6-14 years; each individual underwent 1-3 separate scanning sessions with an interval of approximately one year between each session, resulting in 491 scans) (Fig. 1A). For comparison purposes, we also included cross-sectional rsfMRI data from a group of 61 healthy adults (ages 18-29 years). Both children and adults were scanned using the same scanner and identical scanning protocols. All MR images were subject to a strict quality control process before inclusion in this study (see Supplementary Information for further detail). Using a sliding window approach (window length = 60 s, step size = 2 s), we first derived dynamic functional networks for all rsfMRI scans of each child. Network nodes were defined according to a random parcellation consisting of 1,024 regions of interest of uniform size (Zalesky et al., 2010), and dynamic connections were calculated as the Pearson’s correlation coefficient between the nodal time series within each window. To identify the time-varying modular structure, we employed a multilayer network model (Mucha et al., 2010) by incorporating functional connection patterns from adjacent time windows (Fig. 1B). Then, a temporal modular variability analysis (Liao et al., 2017) was undertaken to quantify how brain nodes spontaneously switched their module affiliations over time (Fig. 1B).

**Figure 1.**
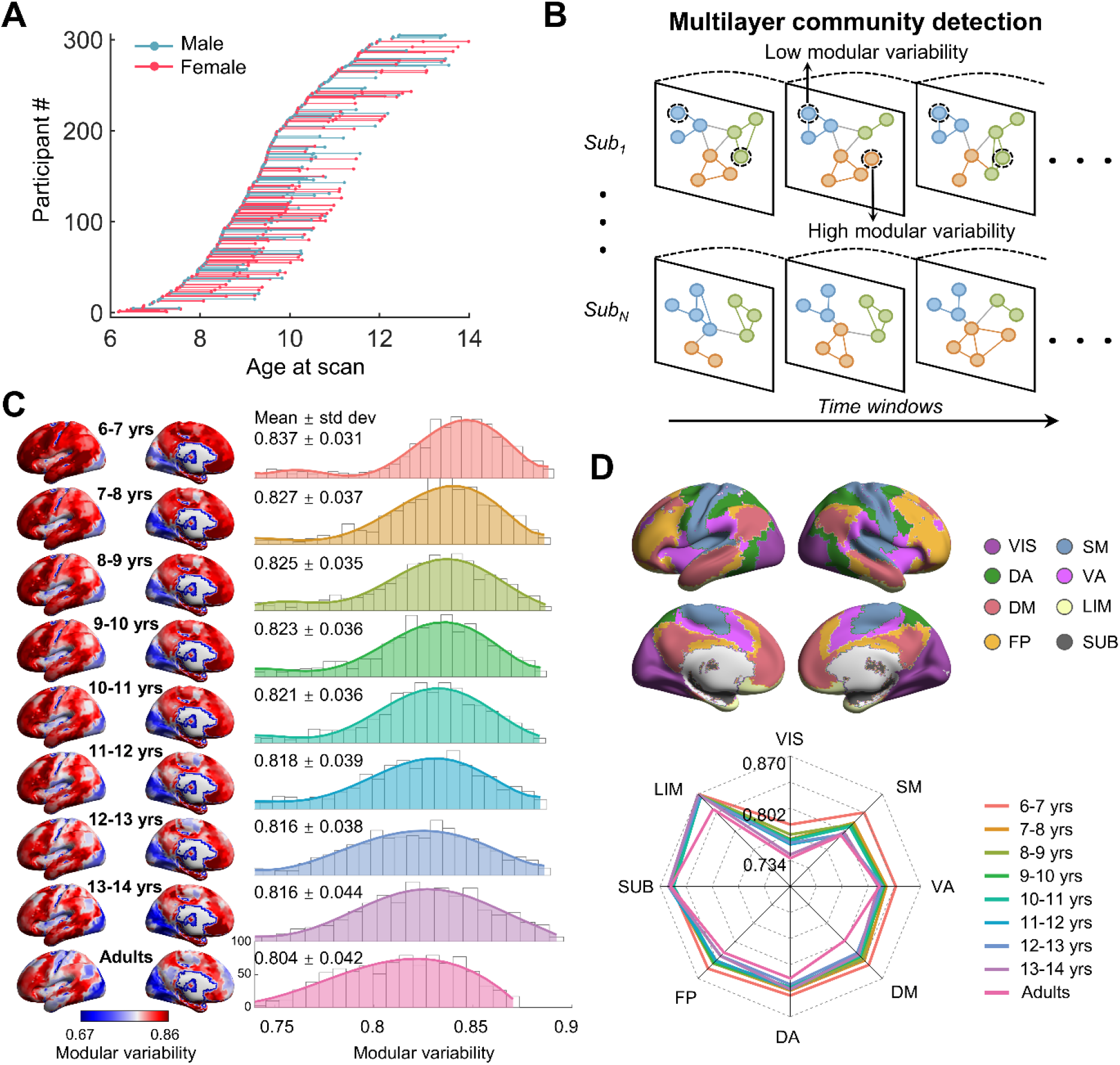
Age distribution of child participants and multilayer module dynamics at different ages. **(A)** Age information of child participant scans. **(B)** Schematic diagram of the multilayer network model and regional modular variability. Each layer represents a functional network within a sliding window. In addition to connections within the same layer, each node also connects to itself in adjacent layers. **(C)** Spatial patterns and frequency polygons of modular variability across the 1,024 nodes for each child subgroup and for the adult group. **(D)** Top: spatial location of the eight functional systems (seven cortical systems (Yeo et al., 2011) and one subcortical system (Tzourio-Mazoyer et al., 2002)). Bottom: distribution of the mean modular variability value of each functional system for each child subgroup and for the adult group. Cortical data was mapped using the BrainNet Viewer software (Xia et al., 2013). VIS, visual; SM, somatomotor; DA, dorsal attention; VA, ventral attention; DM, default-mode; LIM, limbic; FP, frontoparietal; SUB, subcortical.

We found that individual brain networks in children exhibited a modular structure, with modularity (*Q*_*mod*_) values ranging from 0.53 to 0.62 (mean ± std dev = 0.58 ± 0.02) and the number of modules (*N*_*mod*_) ranging from 5.35 to 6.58 (mean ± std dev = 5.93 ± 0.21). To qualitatively illustrate the progressive developmental changes in the spatial pattern of brain dynamics between childhood to adolescence, we divided the 491 rsfMRI scans into eight age subgroups, with a one-year interval between each subgroup. An average modular variability map was then generated for each age subgroup (Fig. 1C). We found that, at a group level, the spatial pattern of modular variability in the child brain network showed regional heterogeneity, with higher variability primarily observed in the frontal and parietal cortices, anterior/middle cingulate gyrus, and middle temporal gyrus, and the lowest modular variability observed in the visual cortex. This spatial pattern observed in the child cohort was highly similar to that of the adult group (Pearson’s correlations: range, 0.83-0.89; mean ± std dev = 0.88 ± 0.02, all *p*_corr_ < 0.0001). The significance levels of these correlations were corrected for spatial autocorrelation using a null distribution of correlation coefficients (Burt et al., 2020) (see Methods section for further detail). When mapping these regions against the eight functional systems (visual, somatomotor, dorsal attention, ventral attention, limbic, frontoparietal, default-mode and subcortical) identified in prior studies (Tzourio-Mazoyer et al., 2002; Yeo et al., 2011), we also observed a corresponding similarity in the system-dependent distributions of modular variability between child and adult brains (Fig. 1D). Notably, the topography of modular variability at both the global and system levels develops in a progressive fashion from childhood through adolescence towards that found in the adult brain. Below, we report our quantitative analysis of the longitudinal development of brain network dynamics.

### Developmental Changes in Brain Network Dynamics in Children

To characterize the longitudinal development of brain network dynamics, a mixed effect model was applied (Diggle & Kenward, 1994; Laird & Ware, 1982). Such models are well suited for cases with missing data, irregular intervals between data measurements, or potential correlation between variables. To account for the potential linear and quadratic age effects, we used two different models that respectively had a linear and quadratic term as their highest-order term, and selected the optimal model with the lower Akaike information criterion value (Akaike, 1974) for use in our analysis (see Methods section for further detail). At the global level, the modularity, *Q*_*mod*_, of the dynamic brain networks increased with age (linear model, *t* = 4.77, *p* < 0.0001, Fig. 2A, left), suggesting enhanced functional segregation of network modules with age. The global mean values of modular variability in the child brain decreased with age (linear model, *t* = -3.28, *p* = 0.0011, Fig. 2A, middle), while the standard deviation across regions increased with age (linear model, *t* = 2.43, *p* = 0.015, Fig. 2A, right). These results suggest that the temporal dynamics of brain networks tend to become more stabilized and more regionally differentiated as the brain develops from childhood to adolescence.

**Figure 2.**
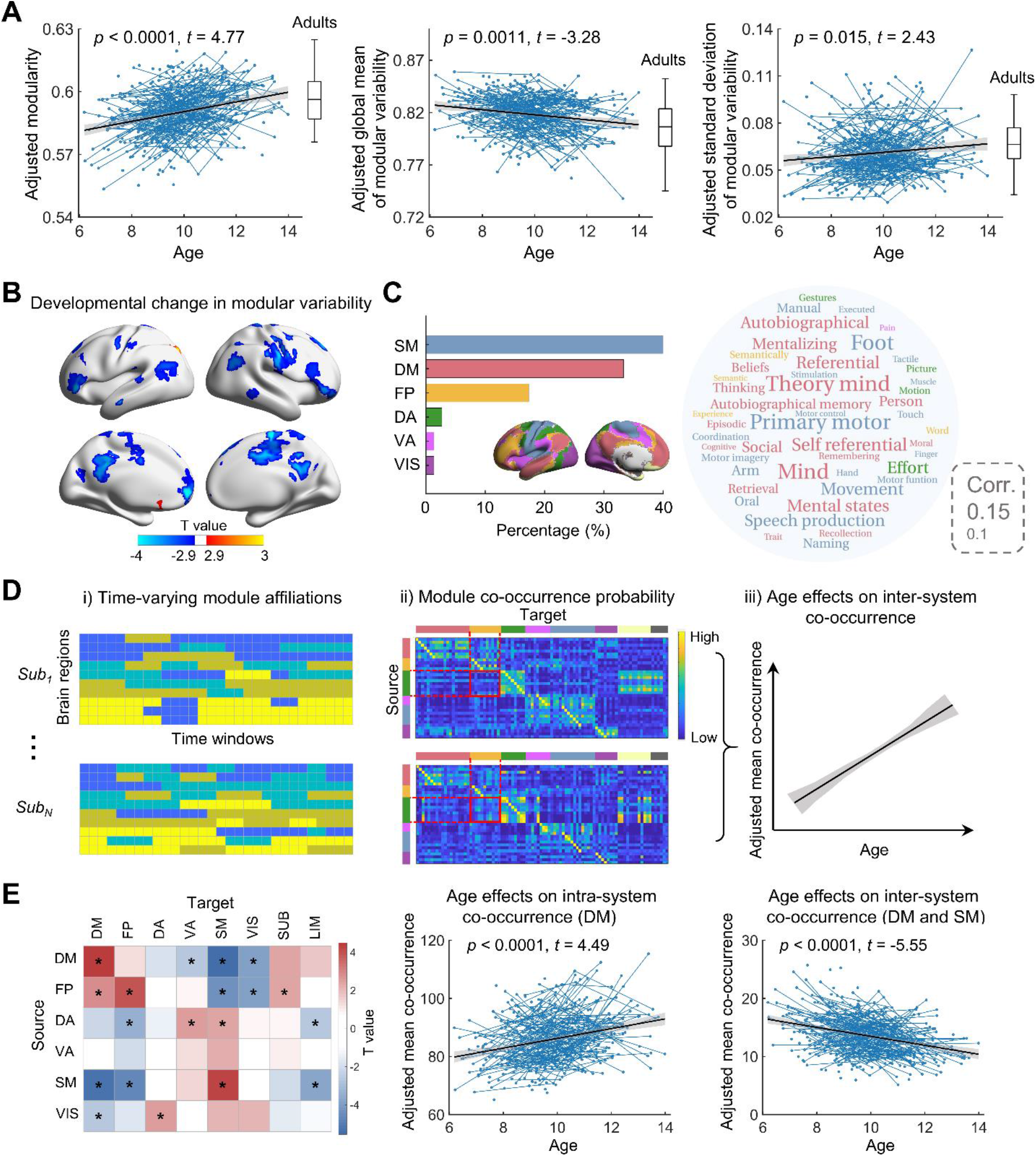
Longitudinal development of brain network dynamics in children. **(A)** Age effects on the network modularity and the mean and standard deviation of the whole brain modular variability. The boxplots represent the distribution median, and the 25th and 75th percentiles of the adult group. **(B)** Spatial distribution of regions showing significant developmental changes in regional modular variability between childhood to adolescence along with their corresponding T values (FDR-corrected *p* < 0.05). **(C)** Left: proportion of regions with significant age-related decreases in modular variability found in each functional system. Right: cognitive terms associated with regions showing significant age-related decreases in modular variability, which was performed based on the NeuroSynth metaanalytic database (Yarkoni et al., 2011). The font size of the cognitive terms represents the value of the correlation coefficient between the regions of interest and the cognitive term maps. Font colors correspond to the different functional systems. **(D)** Schematic diagram of module co-occurrence at the system level. Left: regional module affiliations at each time window, as detected using the multilayer network model. Middle: matrix showing the modular co-occurrence probability between the six source systems and each of the eight functional systems (target systems). Each element in the matrix represents the percentage of time windows in which two nodes from different systems belonged to the same module. Right: estimated age effects on the mean co-occurrence of every system pair. **(E**) Significant developmental effects on modular co-occurrence probability at the system level. Left: age effects on co-occurrence probability (*\*, p* < 0.05, FDR corrected across 48 comparisons at the system level). Middle: age effect on the mean co-occurrence probability for nodes within the default-mode system. Right: age effect on the mean co-occurrence probability of nodes in the default-mode system with those in the somatomotor system. We used a mixed effect model to estimate age effects. In **(A)** and **(E)**, blue lines connecting scattered points represent longitudinal scans of the same child. The adjusted value denotes the measure of interest corrected for sex, head motion, and random age effects. VIS, visual; SM, somatomotor; DA, dorsal attention; VA, ventral attention; DM, default mode; LIM, limbic; FP, frontoparietal; SUB, subcortical.

At the nodal level, we observed that brain regions that showed significant changes in modular variability with age (a total of 77 nodes) predominantly exhibited significant linear decreases in variability (*p* < 0.05, false discovery rate (FDR) corrected). These nodes (75 in total) were principally distributed in the medial and lateral frontal and parietal cortices, supplementary motor area, and somatomotor cortex (Fig. 2B), and were primarily associated with the somatomotor (40%), default-mode (33.33%), and frontoparietal (17.33%) systems. The remaining nodes with significant linear decreases in modular variability with age were spread across the dorsal attention, ventral attention, and visual systems. For ease of reference in our later analysis, we designated these as the six “source systems”. We then explored the cognitive functions associated with these brain regions. Briefly, for each source system, we quantified the spatial correlation between the thresholded t-map denoting significant age effects within the source system and the cognitive term maps available from the NeuroSynth meta-analytic database (Yarkoni et al., 2011). We found that these regions were mainly associated with internal cognitive functions, social inference, and primary motor functions (Fig. 2C). In addition, we also identified one node in the left temporal-occipital junction which showed a significant age-related linear increase, and another node in the left olfactory cortex which showed age-related changes in modular variability that followed a U-shaped quadratic model (*p* < 0.05, FDR corrected).

Next, we examined how the age-related decreases in nodal modular variability (which accounted for the overwhelming majority of age-related changes in variability) were associated with the dynamic interplay between the different functional systems. For each node showing a significant age-related decrease, we computed the modular co-occurrence probability of this node with each of the other nodes by calculating the percentage of time windows during which the two nodes belonged to the same module. Then, the modular co-occurrence of these nodes was summarized at the system level using the functional system atlas derived in previous studies. Specifically, we summed the modular co-occurrence between the six source systems (i.e., the somatomotor, default-mode, frontoparietal, dorsal attention, ventral attention, and visual systems) and all eight functional systems (referred to as the target systems). We assessed age effects on system-level co-occurrence using a mixed effect model with multiple comparison corrections (*p* < 0.05, FDR corrected) (Fig. 2D and 2E).

We observed that, as age increased, nodes in the default-mode, frontoparietal and somatomotor systems showed significantly increased intra-system co-occurrence, indicating enhanced functional specificity within these systems. In relation to inter-system co-occurrence, we found that significant increases in co-occurrence were primarily observed between transmodal areas (consisting of the default-mode/frontoparietal systems) and the subcortical system, as well as between the dorsal/ventral attention systems and the primary sensory systems (i.e., somatomotor and visual systems). Meanwhile, significant decreases in co-occurrence were mainly observed between the default-mode/frontoparietal systems and the primary sensory systems, as well as between the default-mode/frontoparietal system and the attention systems. These findings suggest that the six source systems tend to be divided into two clusters, one comprising the transmodal areas and the subcortical system, and the other comprising the primary sensory and attention systems. During development, functional integration between these two clusters of systems decreases with age.

### Brain Network Dynamics Mediates Age Effects on Communication Efficiency

We further explored whether the development of network dynamics might contribute to age-related changes in the information communication capability of brain networks. Here, we considered two global network metrics, global efficiency (*E*_*glob*_) and local efficiency (*E*_*loc*_) (Latora & Marchiori, 2001; Rubinov & Sporns, 2010) (see Methods section for further detail). *E*_*glob*_ and *E*_*loc*_ respectively capture how efficiently information is transferred across all pairs of nodes, and in the neighborhood of a node. For each scan, the *E*_*glob*_ (or *E*_*loc*_) value of the dynamic brain networks was calculated as the average *E*_*glob*_ (or *E*_*loc*_) value across all windows. We found that *E*_*glob*_ of the functional networks decreased significantly with age (linear model, *t* = -3.34, *p* < 0.001), while *E*_*loc*_ increased significantly with age (linear model, *t* = 4.98, *p* < 0.0001) (Fig. 3A), indicating decreased information integration and increased information segregation between childhood and adolescence. Next, we examined the relationship between brain module dynamics and network efficiency (i.e., *E*_*glob*_ or *E*_*loc*_) across individuals, controlling for age. We found that global mean values of modular variability showed a positive correlation with *E*_*glob*_ (*r* = 0.67, *p* < 0.0001) and a negative correlation with *E*_*loc*_ (*r* = -0.63, *p* < 0.0001) (Fig. 3B). These findings indicate that the dynamic module switching of brain regions is associated with integrated and segregated processing in brain networks.

**Figure 3.**
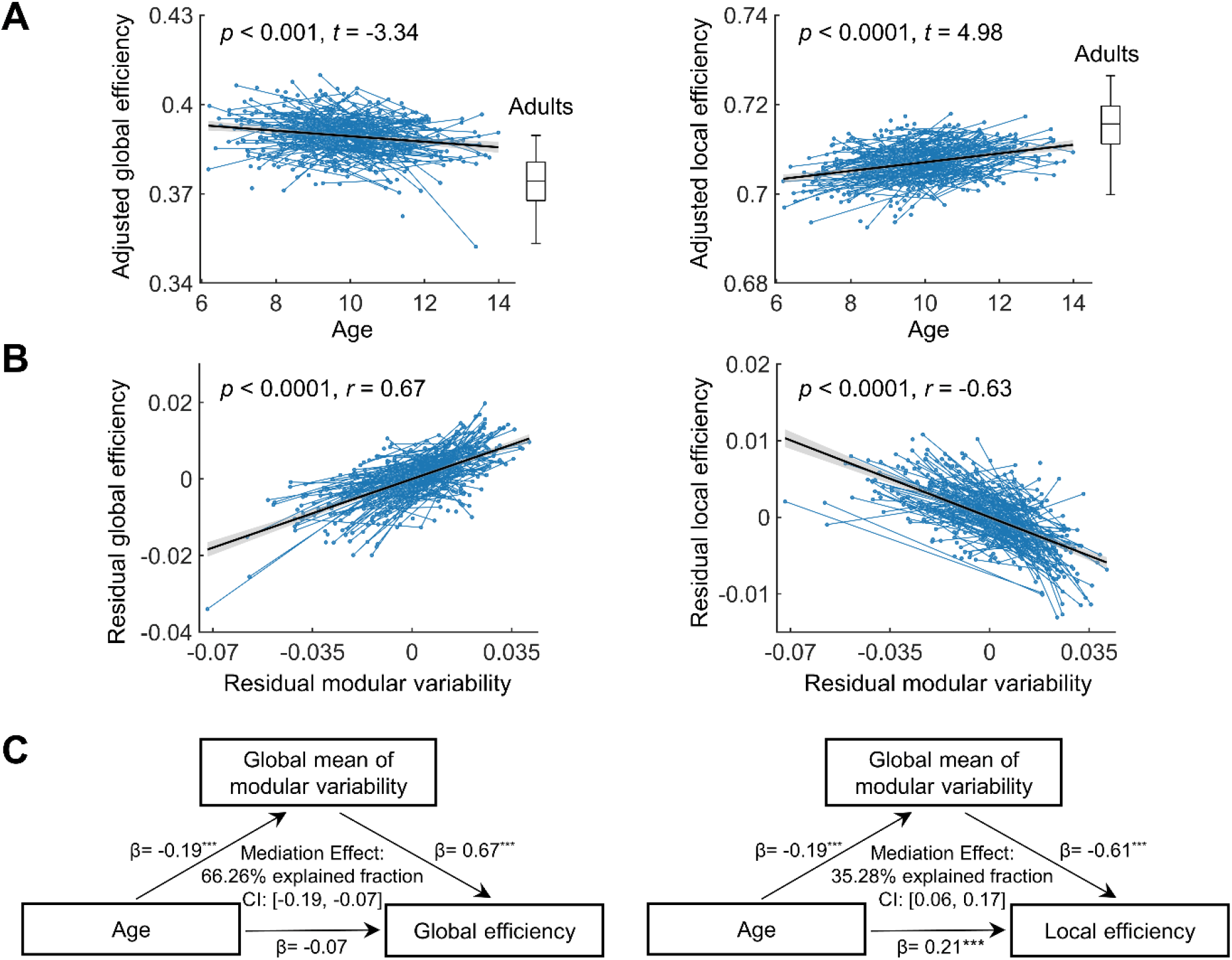
Relationship between age, brain dynamics and network efficiency in children. **(A)** Age effects on global efficiency and local efficiency in the dynamic functional networks. The boxplots represent the distribution median, and the 25th and 75th percentiles of the adult group. **(B)** Relationship between modular variability and network efficiency in children. **(C)** Mediating effects of modular variability on developmental changes in global efficiency (left) and local efficiency (right) (all *p*s < 0.05, bootstrapped n = 5,000). ***, *p* < 0.001. In **(A)** and **(B)**, the blue lines connecting scattered points represent longitudinal scans of the same child. The adjusted value in **(A)** denotes the measure of interest corrected for sex, head motion, and random age effects. The residual value in **(B)** denotes the measure of interest corrected for sex, head motion, and fixed and random age effects. We used a mixed effect model to estimate age effects.

Given the age-related changes in both network efficiency and brain module dynamics, we assessed whether the relationship between age and network efficiency was mediated by brain module dynamics. We performed a mediation analysis (Baron & Kenny, 1986) in which modular variability was taken as the mediator (see Methods section for further detail) and found that the global mean value of modular variability had a significant mediation effect on the relationship between age and *E*_*glob*_ and *E*_*loc*_ (*p* < 0.05 for both, bootstrapped n = 5,000) (Fig. 3C). To further determine the specific brain systems contributing to the mediation effects, we performed a parallel multiple mediation analysis (Preacher & Hayes, 2008), in which the modular variability values of the six source systems (i.e. the somatomotor, default-mode, frontoparietal, dorsal attention, ventral attention, and visual systems) were taken as potential mediators. We found that modular variability of the default-mode, frontoparietal, somatomotor and visual systems exhibited significant mediation effects on the relationship between age and *E*_*glob*_ and *E*_*loc*_ (all *p*s < 0.05, bootstrapped n = 5,000), while modular variability of the attention (i.e., dorsal and ventral attention) system did not (Tables S1 and S2). The explained fraction of the total effect in the mediation models varied across functional systems, being highest in the somatomotor system, followed by the default-mode, visual, and frontoparietal systems for both *E*_*glob*_ and *E*_*loc*_. These findings suggest that the reduction in module dynamics (i.e., network switching) between childhood to adolescence, in particular the reduced dynamics in the somatomotor and default-mode systems, significantly mediates the development of brain network communication efficiency to its mature state in the adult brain.

### Linking Developmental Network Dynamics in Children with Gene Transcriptional Profiles

To investigate whether the regionally heterogeneous development of brain network dynamics was associated with gene expression profiles, we performed a connectome-transcriptome association analysis using brain-wide gene expression data from six donors, publicly available from the Allen Human Brain Atlas (http://human.brain-map.org/) (Hawrylycz et al., 2012). To map the gene expression profiles to the random-1024 parcellation used in our network construction, we first preprocessed the data by performing probe re-annotations, data filtering, probe selection, sample assignment, and data normalization (see Methods section for further details). After preprocessing, expression maps for 15,745 genes were used in the subsequent analysis. Because only data from two donors were available for the right hemisphere, to improve reliability, we conducted our analysis using gene expression data for the left hemisphere, which were available for all six donors. As the majority of nodes showing significant changes in modular variability exhibited age-related decreases (i.e., negative beta values) (Fig. 2B), we explored the spatial association between gene transcriptional profiles and the magnitude of the developmental change (i.e., |*β*_*age*_|) in regional modular variability (Fig. 4A and 4B). For each gene, we estimated the spatial similarity between its transcriptional profile and the absolute value of the developmental rate in modular variability by calculating the Pearson’s correlation coefficient across nodes that showed negative age effects. To correct for potential spatial autocorrelation, we generated a null distribution of correlation coefficients, with the spatial autocorrelation of the original map of developmental rate in modular variability set as a constraint during the generation process (Burt et al., 2020). A total of 4,551 genes were identified as showing a significant correlation (*p* < 0.05, FDR corrected) with developmental changes in modular variability. Of these, 2,190 genes showed a positive correlation and 2,361 genes showed a negative correlation. The 10 genes showing the highest positive correlations are listed in Figure 4C (see Table S3 for all genes showing significant positive correlations). To explore the functional significance of these genes, we performed a gene ontology annotation analysis using the ToppGene Suite (https://toppgene.cchmc.org) (Chen et al., 2009). We found that the positively correlated genes were associated with significant enrichment of biological processes, primarily those involving ion transport and nucleobase-containing compound transport (*p* < 0.05, FDR corrected) (Fig. 4D). For the enrichment of biological processes related to the negatively correlated genes, please see Table S4.

**Figure 4.**
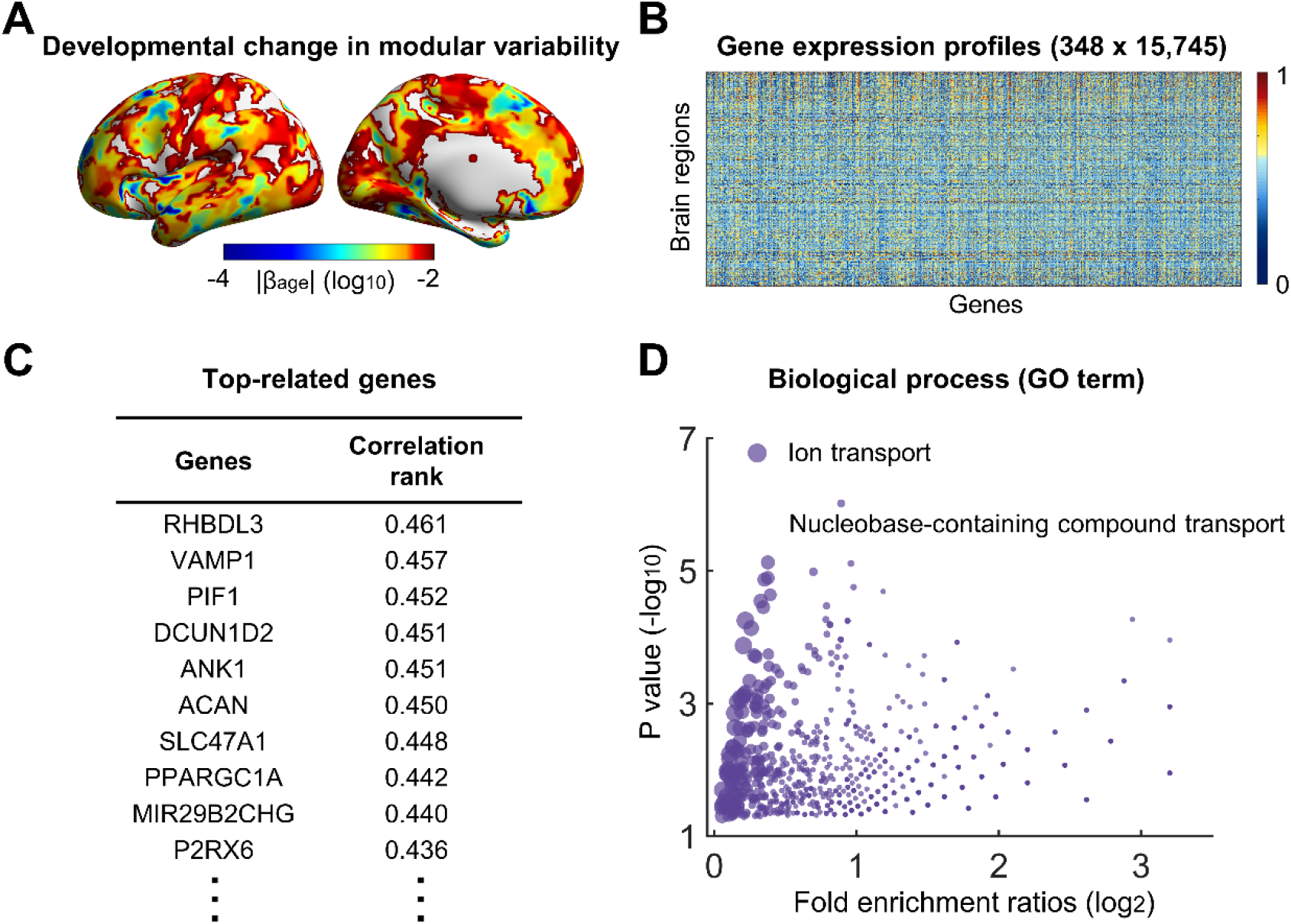
Association between the developmental changes in brain network dynamics and gene expression profiles. **(A)** Spatial pattern of the magnitude of developmental changes (i.e., |*βage*|) in modular variability for regions showing negative linear age effects in the left hemisphere. **(B)** Matrix of gene expression profiles. Each column represents the expression profile of each gene across the 348 nodes of interest. **(C)** Genes showing the highest positive correlations with the developmental change in regional modular variability. Only the top 10 genes are listed. Pearson’s correlation coefficients were calculated within the set of nodes showing negative linear age effects. **(D)** Gene ontology (GO) terms of biological processes associated with genes showing significant positive correlations with developmental changes in module dynamics. Dots marked with text represent GO terms obtained with correction applied for multiple comparisons (FDR-corrected *p* < 0.05), and the remainder represent GO terms obtained where no correction was applied (uncorrected *p* < 0.05). The dot size represents the number of genes that overlap with the corresponding GO term.

### Validations

To demonstrate the robustness of our main findings, we performed several validation analyses, including: (i) testing for the residual effects (post nuisance regression) of head motion on the estimation of modular variability; (ii) evaluating the influence of sliding window parameters on observed brain network dynamics, mediation effects and genetic association by repeating the analyses using a) a longer sliding window length (60 s) and b) weaker coupling between adjacent temporal windows (inter-layer coupling parameter *ω* = 0.75); and (iii) assessing the impact of network thresholding strategies used during functional network construction on observed brain network dynamics, mediation effects and genetic association by repeating relevant analyses on brain networks with an increased network density (density = 10%), and on a different network type (i.e., weighted as opposed to unweighted network).We found that the head motion parameters for mean framewise displacement (FD) (Power et al., 2012) did not show a significant correlation with modular variability (Fig. S1), suggesting that the influence of head motion on the observed developmental changes in brain network dynamics was weak. We also found that, overall, the application of different strategies for analysis did not affect or alter our main conclusions. The results in relation to the development of brain network dynamics, its mediation effects on the network efficiency, and the associated genes remained largely unchanged (Figs. S2-S5).

## Discussion

Using longitudinal rsfMRI data from a large cohort of healthy children, we demonstrate for the first time the development of brain network dynamics between childhood to adolescence and its connection with gene expression profiles. Specifically, modular dynamics in the brain network become progressively more stable as the brain matures, and is correlated with the transcriptional profiles of genes associated with the enrichment of ion transport and nucleobase-containing compound transport. Changes in modular variability occur primarily in the default-mode, frontoparietal and somatomotor systems. The development of more stable network dynamics mediates segregated and integrated processing in the brain, indicating an enhancement in the functional specialization of brain regions during this period of development. Together, these findings highlight the progressive stabilization of network switching between childhood and adolescence and describe the related gene expression profiles, providing insights into the understanding of typical and atypical development. Further, where the majority of previous developmental connectomics studies have mainly focused on dynamic functional connectivity patterns (Faghiri et al., 2018; Hutchison & Morton, 2015; Marusak et al., 2017; Medaglia et al., 2018; Qin et al., 2015; Ryali et al., 2016), our study includes a specific investigation of the maturation of dynamic network topology during childhood and adolescence and the associated cognitive implications, greatly increasing the current knowledge on human brain development.

In the course of our investigation, we also observed that the topography of dynamic modular configurations during childhood and adolescence followed an adult-like spatial pattern. Evidence from adults indicates that during rest, brain regions spontaneously switch between functional modules in a spatially heterogeneous way, such that the association cortex shows higher temporal variability than the primary cortex (Chen et al., 2016; Liao et al., 2017; Liu et al., 2020; Pedersen et al., 2018). Highly variable regions usually act as flexible hubs to maintain efficient inter-module communication and promote cognitive flexibility (Schaefer et al., 2014; Yin et al., 2020). The spatial pattern observed from our child rsfMRI dataset showed a regional distribution of modular variability similar to that of the adult brain described in previous studies. A similar spatial pattern was also observed in a previous study on functional module switching in infants (Yin et al., 2020). Combining our observations with prior findings in studies on the adult (Chen et al., 2016; Liao et al., 2017; Liu et al., 2020; Pedersen et al., 2018) and infant brain (Yin et al., 2020), we speculate that the spatially heterogeneous pattern is a common and underlying property that reflects the regional diversity in brain module dynamics.

The age-related decrease in functional modular variability may be explained by the development of white matter structural connections. Temporal variability in functional connectivity strength has been demonstrated to be structurally constrained by white matter tracts (Deco et al., 2011; Fukushima et al., 2018; Liao et al., 2015; Zhang et al., 2016). More specifically, two brain regions linked by direct structural connections (i.e., white matter tracts) tend to show smaller temporal variability in functional connectivity strength than regions without direct structural connections, and the greater the strength of a direct structural connection, the smaller the temporal variability of the functional connectivity strength between the regions (Liao et al., 2015). Between childhood to adolescence, white matter tracts undergo profound refinements, including regressive processes (e.g., elimination of circuits, axonal projections or synapses) and progressive myelination at the microscopic scale (Tau & Peterson, 2010; Vértes & Bullmore, 2015), and heterogeneous increases in structural connectivity strength at the macroscopic scale (Huang et al., 2015; Zhao et al., 2015). Given that structure-function coupling increases with age (Baum et al., 2020; van den Heuvel et al., 2015), the development of the white matter structural network may reduce regional switching frequency between functional modules and thereby promote the development of modular dynamics in the brain towards an adult level of stability.

Enhanced functional segregation during development has been identified in previous neurodevelopmental studies which applied a static functional connectivity approach (Cao et al., 2016; Gu et al., 2015; Stevens et al., 2009). This is further confirmed by our findings that functional modularity in dynamic networks also increases with age. Consistent with the developmental changes in functional modularity, we also observed that, between childhood and adolescence, global network efficiency decreased and local network efficiency increased with age. Interestingly, we found that regional modular variability, especially that of the default-mode and somatomotor systems, significantly mediated the relationship between age and network efficiency. Prior studies in adults have demonstrated that dynamic adjustments in connectivity, especially connectivity adjustments in the default-mode network, induce fluctuations in network efficiency over time (de Pasquale et al., 2016; Zalesky et al., 2014). Thus, it is reasonable to postulate that, between childhood to adolescence, reduced modular variability in the default-mode system contributes to changes in intra- and inter-module information communication, affecting the development of communication efficiency in the dynamic functional networks.

In developmental cognitive neuroscience, the theory of interactive specialization (Johnson, 2011) posits that during postnatal development, the function of brain regions becomes more specialized, as a result of the age-related reconfiguration of inter-regional interactions driven by intrinsic activities or environmental stimuli. Considering that brain module dynamics plays a crucial role in individual cognition (e.g., working memory) and behavior (e.g., motor skill learning) in adults (Bassett et al., 2011; Braun et al., 2015; Shine & Poldrack, 2018), the reduction in regional module variability and the changes in network efficiency observed in our study may be related to individual refinements in these capabilities during childhood and adolescence. Consistent with this, we found that the regions showing most significant decreases in modular variability were those primarily involved in self-referential thinking, social cognition and motor functions. In addition, a previous study has suggested that network flexibility in the human brain decreases when turning a motor skill task into an automatic process after a period of training (Bassett et al., 2015). We found that segregation between the somatomotor system and high-order systems increased between childhood to adolescence, suggesting that the decrease in modular variability in this system could be related to the refinement of somatomotor capabilities during this period. However, in contrast to our observations, one recent study found that regional module switching (i.e., flexibility), especially that of the primary system, showed significant increases with age during the first two years of life (Yin et al., 2020). This discrepancy may be attributable to the different developmental phases considered (infants versus school-age children) or to the application of different network construction strategies (i.e., absolute correlation thresholding versus fixed-density thresholding) in the two studies.

By performing a connectome-transcriptome association analysis, a recent study that we also undertook demonstrated that the spatial heterogeneity of module dynamics in the adult brain is shaped by the expression profile of the genes primarily associated with potassium ion transport (Liu et al., 2020). Nevertheless, the genetic basis underpinning the development of functional network dynamics remains poorly understood. Our work addresses this gap in knowledge by revealing that the maturation of brain module dynamics towards an adult-like state is associated with the expression profiles of genes associated with the enrichment of ion transport and nucleobase-containing compound transport. Ion transport is one of the most important functions of a neuron, facilitating the balance of ion concentrations in and out of the cell membrane and promoting the stability of brain neural circuits (Gibson et al., 2007). A recent computational modeling study further suggests that ion concentration dynamics causes spontaneous neuronal fluctuations (Krishnan et al., 2018), which may contribute to fluctuations in blood oxygenation level-dependent (BOLD)-fMRI signals (Schölvinck et al., 2010). In addition, nucleoside transport has been found to be dependent on ion concentrations (especially that of Na^+^), and Na+/nucleoside co-transports are also an electrogenic process (Griffith & Jarvis, 1996). Given the above, it makes sense that the development of adult-like modular dynamics is related to the transport of ions and nucleobase-containing compounds, which can affect inter-regional interactions by modulating neural activities.

Of the genes that were found in our study to be highly correlated with age related decreases in modular variability, several have also been described in prior studies as being related to brain development. Specifically, ACAN (aggrecan) has been found to control the maturation of glial cells during brain development (Dornowicz et al., 2008); the basic helix-loop-helix gene HES6 promotes neuronal differentiation (Bae et al., 2000); FGF9 (fibroblast growth factor 9) is crucial for the postnatal migration of cerebellar granule neurons (Lin et al., 2009); and Sema7A (semaphorins) regulates climbing fiber synapse elimination in the developing mouse brain (Uesaka et al., 2014). The identification of correlated genes provides novel clues for bridging the gap between our understanding of the developmental changes in brain network dynamics and our limited knowledge of the underlying biological mechanisms behind this process.

Several issues and future research topics also come to light for further consideration. First, postmortem gene expression data from adult donors obtained from the Allen Human Brain Atlas was used to explore the relationship between gene expression and the age-related changes of network dynamics in children. However, while the absolute expression levels of genes may change with age due to developmental effects, their spatial patterns do not seem to change greatly after birth (Kang et al., 2011). As our interest lies in the spatial pattern (i.e., relative values across regions) of gene expression profiles rather than exact expression values, the choice of gene expression data should not have a great influence on our findings. Nevertheless, the availability of cerebral cortex gene expression data for children and adolescents could be beneficial for facilitating future exploration of the molecular mechanisms underlying developmental network dynamics. Secondly, previous studies in adults have suggested that brain network dynamics show a relationship with cognitive flexibility (Chen et al., 2016; Liao et al., 2017; Yin et al., 2020) and individual task performance (Pedersen et al., 2018). How then is the progressive maturation of brain network dynamics during childhood and adolescence associated with the development of individual cognition and behavior? This still remains to be elucidated. Finally, as abnormalities in the dynamic characteristics of functional brain networks have been observed in several neurodevelopmental disorders (e.g., attention-deficit/hyperactivity disorder (Ding et al., 2020) and autism spectrum disorder (Harlalka et al., 2019)), delineating the typical developmental trajectory of brain network dynamics may provide novel clues for the early detection or diagnosis of atypical neurodevelopment.

## Methods

### Participants

We utilized a longitudinal rsfMRI dataset consisting of scans obtained from 360 typically developing children (F/M = 163/197, 6 to 14 years, 643 scans in total) collected by the Children School Functions and Brain Development Project (Beijing Cohort). Participants included in this study were cognitively normal, and had no history of neuropsychiatric illness, psychoactive drug use, significant head injuries, or significant physical illness. Some of these children underwent multiple sessions of multi-modal MRI imaging (T1, T2, rsfMRI, etc.) with an interval of approximately one year between each session. After strict quality control screening (see Supplementary Information for further detail), 491 rsfMRI scans of 305 children (F/M = 143/162, 6 to 14 years) were retained for use in our study. These were made up of 3 scans from 47 children (F/M = 31/16), 2 scans from 92 children (F/M = 47/45), and a single scan from 166 children (F/M = 65/101). For the purposes of comparison, we also made use of an rsfMRI dataset comprised of data from 61 healthy young adults (F/M = 37/24, 18 to 29 years), which was acquired using an identical scanner and scanning protocols. The study was approved by the Ethics Committee of Beijing Normal University, and written informed consent was obtained from all participants or their parents/guardians.

### Data acquisition

MRI data were acquired using a 3T SIEMENS Prisma scanner in the Center for Magnetic Resonance Imaging Research at Peking University. For each participant (child and adult), structural and functional MRI scans were acquired using the following protocols. T1-weighted images were acquired using a sagittal 3D magnetization prepared rapid acquisition gradient echo (MPRAGE) sequence: repetition time (TR) = 2,530 ms, echo time (TE) = 2.98 ms, inversion time = 1,100 ms, flip angle (FA) = 7°, matrix = 256 × 224, field of view (FOV) = 256 × 224 mm^2^, slice number = 192, slice thickness = 1 mm, bandwidth = 240 Hz/Px. The rsfMRI data was acquired using an echo-planar imaging sequence: TR = 2,000 ms, TE = 30 ms, FA = 90°, matrix = 64 × 64, FOV = 224 × 224 mm^2^, slice number = 33, slice thickness/gap = 3.5/0.7 mm, scan duration = 8 minutes (i.e., 240 volumes in total). The participants were asked to fix their vision on a bright cross-hair in the center of the scanner screen. A field map was acquired prior to the rsfMRI scan using a 2D dual gradient-echo sequence: TR = 400 ms, TE1 = 4.92 ms, TE2 = 7.38 ms, FA = 60°, matrix = 64 × 64, FOV = 224 × 224 mm^2^, slice number = 33, slice thickness/gap = 3.5/0.7 mm.

### Data preprocessing

Resting state fMRI data from the child cohort was preprocessed using SPM12 (https://www.fil.ion.ucl.ac.uk/spm) and DPABI 3.0 (Yan et al., 2016). First, for each scan, we removed the first ten volumes and performed slice-timing correction. Next, a field map correction was applied to remove geometric distortion. We then performed head motion correction and estimated the mean FD (Power et al., 2012) across time for each scan. Ninety-four scans were excluded due to excessive head motion (i.e., translation > 3 mm, rotation > 3°, or mean FD > 0.5 mm). The functional images were then co-registered with individual T1 images and spatially normalized to a custom template using a unified segmentation algorithm (Ashburner & Friston, 2005) (see Supplementary Information for further detail). During initial segmentation of the T1 images, Chinese Pediatric Atlases (CHN-PD) (6-12 years) (Zhao et al., 2019) were used as the reference for segmentation to improve accuracy in the spatial deformation of pediatric brain images. The normalized functional images were resampled to 3-mm isotropic voxels and spatially smoothed with a Gaussian smoothing kernel (full-width at half maximum = 4 mm). Next, we performed linear detrending, nuisance signal regression, and temporal band-pass filtering (0.01-0.1 Hz). During nuisance regression, the following nuisance regressors were included as covariates to reduce the influence of non-neural signals: Friston’s 24 head motion parameters (Friston et al., 1996), “bad” time points with FD above 0.5 mm, and white matter, cerebrospinal fluid and global brain signals.

Functional images from the adult cohort were preprocessed using the same procedures, except that when undertaking spatial normalization, functional images from the adult group were spatially normalized to the Montreal Neurological Institute (MNI) standard space.

### Construction of dynamic functional networks

The selection of an appropriate brain parcellation scheme is essential for node definition in the construction of functional networks (Bullmore & Bassett, 2011). For functional networks constructed from the child scans, we defined network nodes based on a customized random parcellation scheme, referred to as random-1024 parcellation, comprising 1,024 gray matter regions of uniform size (Zalesky et al., 2010). The time course for each node was extracted by averaging the time courses across voxels within the node. We then applied a commonly used sliding window approach (window length = 60 s, step size = 1 TR (i.e., 2 s), total windows for each scan = 201) to estimate dynamic functional connectivity over time (Hutchison et al., 2013; Lurie et al., 2020). Inter-node functional correlations for each window were approximated using the Pearson’s correlation coefficient between nodal time courses. The resulting networks were then thresholded by applying a network density of 5% to remove weak or spurious connections introduced by noise, producing a time-varying binary functional network for each rsfMRI scan of each child. Negative correlations were eliminated prior to network thresholding due to their ambiguous physiological interpretation (Fox et al., 2009; Murphy & Fox, 2017).

Functional networks from the adult scans were constructed using the same procedures. To enable regional-level comparisons between the child and adult networks, we obtained the parcellation scheme for the adult networks by spatially transforming the random-1024 parcellation from the children’s custom space to MNI space.

### Identification of dynamic modular architecture

We employed a multilayer network model (Mucha et al., 2010), which can incorporate connectivity information within adjacent time windows, to map the dynamic modular structure in the child and adult brain networks. Specifically, the dynamic functional networks in each scan were considered as a multilayer network consisting of 201 time-ordered layers (i.e., windows) with ordinal inter-layer coupling, in which identical nodes in adjacent layers were coupled with nonzero strength. Then, we identified the time-dependent modular architecture by optimizing the modularity, *Q*_*mod*_, of the multilayer network, with an implicit assumption that module change between layers was continuous. The modularity, *Q*_*mod*_, of the time-varying modular structure is defined as (Mucha et al., 2010):

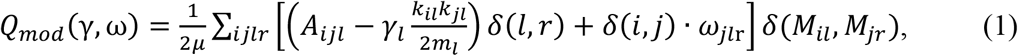

where variables *i* and *j* are node labels, and variables *l* and *r* are layer labels. Specifically, *μ* represents the total connectivity strength of the entire network, *m*_*l*_ denotes the total connectivity strength within layer *l, A*_*ijl*_ denotes the connectivity strength between node *i* and node *j* in layer *l, k*_*il*_*k*_*jl*_/2*m*_*l*_ denotes the connection probability expected by chance between node *i* and node *j* in layer *l, k*_*il*_ denotes the degree of node *i* in layer *l*, and *M*_*il*_ denotes the module label of node *i* in layer *l*. The function *δ* (*x, y*) is equal to 1 if variable *x* is identical to variable *y*, and is equal to 0 in all other cases. Parameters *γ* and *ω* are, respectively, the topological resolution parameter and temporal coupling parameter. Parameter *γ* defines the module size. The larger the value of *γ*, the smaller the size of the identified modules, and the greater the number of modules in the network. Here, we used the commonly-used default value of *γ* = 1 (Bassett et al., 2011; Braun et al., 2015). Parameter *ω* determines the extent of inter-layer interaction. The smaller the value of *ω*, the more independent the adjacent layers. We chose *ω* = 1 to balance the influence of inter-layer and intra-layer edges (when *ω* <1, intra-layer edge strength dominates modularity optimization) (Bassett et al., 2011; Braun et al., 2015). Notably, the dynamic modular architecture varies slightly with each instance of mapping since the heuristic Louvain algorithm (Blondel et al., 2008) is applied in the optimization of modularity. Here, all measurements relating to dynamic modular architecture were taken as the average across 100 instances of mapping. The module detection algorithm for multilayer networks was obtained from an open MATLAB code package at http://netwiki.amath.unc.edu/GenLouvain/GenLouvain (Jeub et al., 2012).

### Modular variability analysis

To characterize the temporal reconfiguration of functional modular architecture, we tracked the change in functional modules over each window. Specifically, we assessed the change in module affiliation (i.e., network switching) over time for all nodes using modular variability as the chosen metric (Liao et al., 2017). First, given a node *i*, we evaluated the variability of its module affiliation between any two windows *t* and *t’* (Steen et al., 2011) as

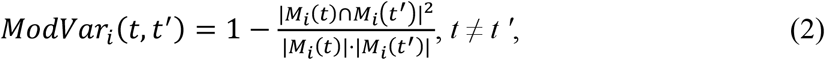

where M_*i*_(*t*) denotes the module to which node *i* belonged in window *t*, |M_*i*_(*t*)| represents the number of nodes included in module M_*i*_(*t*), and |M_*i*_(*t*)∩M_*i*_(*t’*)| represents the number of nodes in the intersection between modules M_*i*_(*t*) and M_*i*_(*t’*). A small intersection between two modules indicates large variability. Secondly, we calculated the total modular variability of a node over all time windows as (Liao et al., 2017)

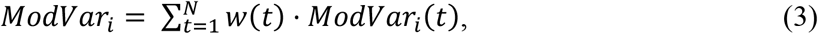

where 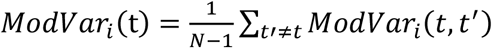 denotes the modular variability of node *I* between window *t* and all other windows, and *N* denotes the total number of windows. A normalized weighted coefficient *w*(*t*) is employed in Eq. (3) to reduce the impact of potential outlier windows. The coefficient *w*(*t*) is a measure of inter-window similarity, which is calculated using adjusted mutual information (Vinh et al., 2010), and denotes the overall similarity of the modular structure in window *t* to that in all other windows. For each rsfMRI scan, we calculated the modular variability of each of the 1,024 nodes. The larger the nodal modular variability, the more often the node tends to switch between modules over time. The mean modular variability of the whole brain was calculated as the average modular variability across all nodes.

To illustrate the change in modular variability patterns with development, we divided all child rsfMRI scans into eight subgroups, with a one-year interval between each subgroup. A group-level modular variability map was generated for each subgroup by averaging individual maps within each subgroup. To enable comparison, a group-level modular variability map for the young adults was also generated. We then conducted Pearson’s correlation analyses to measure the spatial similarity between the modular variability maps of each age-group (i.e., all child subgroups versus the adult group). To correct for spatial autocorrelation, we generated 10,000 surrogate maps constrained by the spatial autocorrelation characteristics of the modular variability map generated for the adult group (Burt et al., 2020), and obtained a null distribution of correlation coefficients for each subgroup of children. Empirically observed spatial similarity values were compared against the null distribution to determine significance levels. To further assess the system dependence of nodal modular variability, we categorized the 1,024 nodes into eight functional systems. Seven of these systems were obtained from a prior functional system parcellation scheme (Yeo et al., 2011): the visual, somatomotor, dorsal attention, ventral attention, limbic, frontoparietal, and default-mode systems. The remaining subcortical system was extracted from the Automated Anatomical Labeling atlas (Tzourio-Mazoyer et al., 2002). When analyzing system-dependence in children, the functional system atlases defined from adults were spatially transformed to the children’s custom space prior to the allocation of nodes to their respective functional systems. Finally, we quantified the age-related changes in regional modular variability using a mixed effect model.

### Relationship between developmental changes in brain network dynamics and cognitive function

We explored the cognitive significance of regions showing significant age-related changes in modular variability using the NeuroSynth meta-analytic database (www.neurosynth.org) (Yarkoni et al., 2011). Specifically, we examined the cognitive terms associated with the regions exhibiting significant age-related decreases in modular variability, since these regions made up the overwhelming majority of brain areas showing developmental changes in network dynamics. We first generated six thresholded *t*-maps denoting age effects on regions of interest within each functional system separately by mapping these regions into the six source systems (i.e. the somatomotor, default-mode, frontoparietal, dorsal attention, ventral attention, and visual systems) (Tzourio-Mazoyer et al., 2002; Yeo et al., 2011). Next, we quantified the Pearson’s correlation between each *t*-map and all cognitive term maps available from the NeuroSynth database. The results were illustrated using word-cloud plots.

### Module co-occurrence analysis at the system level

To explore whether the developmental changes in nodal modular variability were associated with the dynamic interplay between functional systems, we examined the age-related changes in module co-occurrence of different systems. Since most nodes exhibiting significant age effects showed linear decreases with age (Fig. 2B), we focused our analysis on nodes showing a significant negative age effect (*N*_*s*_ nodes in total). Briefly, the module co-occurrence probability of each node showing significant age effects with each of the other nodes in the brain network was calculated as the percentage of time windows in which the two nodes belonged to the same module (Braun et al., 2015). For all scans of each child, an *N*_*s*_ × 1,024 co-occurrence matrix was obtained (Fig. 2D). Next, the module co-occurrence matrix was summarized at the system level by calculating the average co-occurrence probability of nodes in each source system with nodes in each of the respective target systems. (Source systems are those containing nodes showing significant age-related decreases, i.e., the somatomotor, default-mode, frontoparietal, dorsal attention, ventral attention, and visual systems. Target systems refer to all eight functional systems.) Therefore, for all scans of each child, we obtained a 6 × 8 module co-occurrence matrix at the system level, each row of which denoted the co-occurrence probability of a source system with each of the target systems. Finally, we assessed the age-related changes in the co-occurrence probability for each pair of systems using a mixed effect model. The significance level was corrected for multiple (i.e., 48) comparisons at the system level using the FDR method (corrected *p* < 0.05).

### Relationship between age, brain dynamics and network efficiency

To explore whether dynamic modular reconfiguration was related to information communication capabilities in the brain network, we considered two global network metrics, global efficiency and local efficiency, to capture different aspects of information transmission efficiency.

#### Global efficiency (*E*_*glob*_)

Global efficiency measures information transmission efficiency across all pairs of nodes in the network (Latora & Marchiori, 2001; Rubinov & Sporns, 2010). In a given network, *E*_*glob*_ is defined as

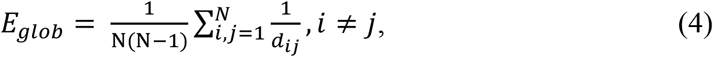

where *N* represents the total number of nodes in the network, and *d*_*ij*_ is the shortest path length between node *i* and node *j. E*_*glob*_ of the dynamic brain network was calculated as the average *E*_*glob*_ across all time windows.

#### Local efficiency (*E*_*loc*_)

Local efficiency measures information communication efficiency between local subgraphs (Latora & Marchiori, 2001; Rubinov & Sporns, 2010). In a given network, *E*_*loc*_ is defined as

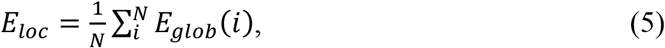

where *N* represents the total number of nodes in the network, and *E*_*glob*_(*i*) represents the *E*_*glob*_ of the neighborhood nodes of node *i. E*_*loc*_ of the dynamic network was calculated as the average *E*_*loc*_ across all time windows.

We estimated both the *E*_*glob*_ and *E*_*loc*_ of the dynamic functional networks for every scan, and applied a mixed effect model to explore age effects on these two measures. To assess whether network efficiency was related to modular variability, we conducted a Pearson’s correlation analysis between the mean modular variability of the brain and network efficiency across all scans, correcting for age, sex, and head motion effects. To further explore whether brain module dynamics mediated the age effects on network efficiency, we performed a single-level mediation analysis, with age, the mean modular variability of the brain, and network efficiency (i.e., *E*_*glob*_ and *E*_*loc*_), set respectively as the independent variable (X), mediator (M) and dependent variable (Y). Finally, to differentiate the contribution of different functional systems to the mediation effect, we employed a parallel multiple mediation analysis, with the mean modular variability of each of the six source systems (i.e. the somatomotor, default-mode, frontoparietal, dorsal attention, ventral attention, and visual systems) designated as mediator. In each mediation model, the explained fraction of the total effect for a given indirect path was defined as the product of the standard regression coefficients along this path divided by the sum of the products for all paths. The mediation analysis was performed using the PROCESS plugin in SPSS (Preacher & Hayes, 2008). We then undertook bootstrapping (n = 5,000) to assess the statistical significance of the mediation analysis, for which a 95% confidence interval without 0 was equivalent to a significance level of 0.05 (Preacher & Hayes, 2004).

### Relationship between developmental changes in brain network dynamics and gene expression profiles

To investigate the association between developmental changes in brain network dynamics and gene expression profiles, we used brain-wide gene expression data publicly available from the Allen Human Brain Atlas (http://human.brain-map.org/) (Hawrylycz et al., 2012). This atlas contains 3,702 tissue samples from six donors, and provides their accurate MNI coordinates. Samples from two donors cover the whole brain, and the samples from the remaining four donors cover only the left hemisphere. Using the minimally processed data provided in the Allen Human Brain Atlas (http://help.brainmap.org/display/humanbrain/Documentation), we carried out the following procedures. First, we removed samples located in the brain stem and cerebellum, and re-annotated the gene names of probes for the remaining 2,748 samples. Secondly, we used the intensity-based filtering method (Arnatkevičiūtė et al., 2019) to filter the data. For each gene, its expression level in a given sample was obtained by averaging the expression values across all detecting probes. Next, we normalized the expression data using the scaled robust sigmoid (SRS) algorithm (Fulcher et al., 2013) (see Supplementary Information for further detail). We then matched the MNI coordinates of each sample to the random-1024 parcellation scheme of the adult group using the nearest-point search algorithm. Each sample was then assigned to one of the brain nodes. For each node, expression data for each gene was obtained by first averaging the data across samples from the same donor, and then averaging the nodal expression data across donors. Using this process, for each node, we obtained gene expression data for 15,745 genes. The preprocessing of gene expression data described above was performed by referencing the code at https://github.com/BMHLab/AHBAprocessing (Arnatkevičiūtė et al., 2019). As gene expression data for the right hemisphere was available from only two donors, to improve the reliability, we used left hemisphere gene expression data from all six donors for the subsequent brain network-gene association analysis.

To examine whether the spatial inhomogeneity in developmental changes in nodal modular variability was associated with gene expression levels, we performed a spatial similarity analysis across the 348 nodes showing negative linear age effects (i.e., age-related *β*_*age*_ < 0). For each of the 15,745 genes, we undertook a Pearson’s correlation analyses to measure the spatial similarity between gene expression profile and the magnitude of developmental changes (i.e., |*β*_*age*_|) in modular variability. To correct for spatial autocorrelation, we generated 10,000 surrogate maps constrained by the spatial autocorrelation characteristics of the map of developmental changes in modular variability (Burt et al., 2020), and obtained a null distribution of correlation coefficients for each gene of interest. Empirically observed spatial similarity values were compared against the null distribution to determine significance levels. Significantly correlated genes were identified with an FDR-corrected *p* < 0.05 and further divided into two categories, i.e., genes showing a significant positive correlation with age-related changes in modular variability and those showing a significant negative correlation. Separate functional enrichment analyses on these two categories of genes were performed using the ToppGene Suite (https://toppgene.cchmc.org/) (Chen et al., 2009).

### Statistical modeling

Given the nature of the longitudinal rsfMRI data used in this study, we applied a mixed effect model to study the developmental trajectories of brain network measures (Diggle & Kenward, 1994; Laird & Ware, 1982). Such models are well suited for cases with missing data, irregular intervals between data measurements, or potential correlation between variables. To account for potential linear and quadratic age effects, we undertook separate analyses using two different models that respectively had a linear term and quadratic term as their highest-order term. For each analysis, we used the maximum likelihood method to undertake parameter estimation, and applied the Akaike information criterion (Akaike, 1974) to select the optimal model. Specifically, the linear model was defined as

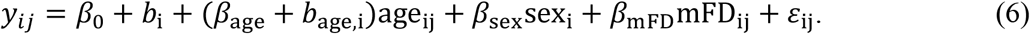

The quadratic model was defined as

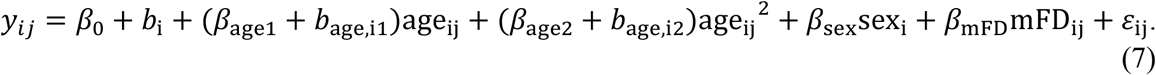

In these models, *y*_*ij*_ represents the observed brain network measures of subject *i* at the *j*th scan, *β*_*age*_ represents the fixed age effects, *b*_*age,i*_ represents the random effects of subject *i*, and represents the residual of subject *i* at the *j*th scan. Sex and mean FD (mFD) were also included as covariates in the two models. Here, we used the statistical models to estimate age effects on the following measures: the modularity (*Q*_*mod*_) of the dynamic networks, the mean and standard deviation of regional modular variability across the brain, nodal modular variability values, and network efficiency (*E*_*glob*_ and *E*_*loc*_). All scatter plots illustrate fixed age effects after correction for random effects.

### Validation Analyses

We further investigated whether our results were affected by head motion and network analysis strategies, specifically the choice of sliding window length, temporal coupling strength between adjacent windows, and network thresholding strategies. First, previous studies suggest that head motion can introduce spatially inhomogeneous bias in the estimation of functional connectivity (Power et al., 2012; Power et al., 2015), which may affect the evaluation of developmental effects (Satterthwaite et al., 2013). To reduce the influence of head motion, we included Friston’s 24 head motion parameters (Friston et al., 1996), the global brain signal, and “bad” time points (FD > 0.5 mm) as covariates during nuisance regression. To further assess the residual influence of head motion, we also calculated the Pearson’s correlation coefficient between head motion (i.e., mean FD) and global network dynamics across scans. Secondly, the selection of sliding window length affects the estimation of dynamic connectivity and thus the temporal characteristics of the functional networks (Hutchison et al., 2013; Lurie et al., 2020; Shakil et al., 2016). In our main analysis, we set the window length as 60 s, a timeframe which is able to reliably capture the temporal variations in the functional networks (Zalesky et al., 2014). To assess the potential influence of the sliding window length on our findings, we reconstructed the dynamic networks using a window length of 100 s. Third, in the multilayer network model, the temporal coupling parameter, *ω*, between adjacent windows has a strong impact on the reconfiguration of modular architecture between windows (Mucha et al., 2010). In addition to using *ω* = 1 in our main analysis, we also repeated the multilayer network module mapping analysis using *ω* = 0.75. Finally, we evaluated whether our findings were influenced by the network thresholding strategies applied in functional network construction, which can affect the estimation of the graph metrics (Bullmore & Bassett, 2011). In our main analysis, the maturation of network topology was our principal focus, and we therefore generated binary networks with a fixed network density (i.e., 5%) to correct for inter-subject differences in the number and strength of functional connectivities. To explore the influence of network density, we constructed binary networks with a density of 10%. In addition, we also constructed weighted functional networks with a density of 5% to assess the impact of connectivity strength.

## Data availability

Nodal time series of preprocessed rsfMRI signals and some data supporting the results of this study are available at https://github.com/helab207/Development-of-brain-module-dynamics.

## Code availability

Codes used for the neuroimaging analysis and the statistical models are available at https://github.com/helab207/Development-of-brain-module-dynamics.

## Acknowledgments

The study was supported by the National Natural Science Foundation of China (Nos. 82021004, 81971690, 31830034, 81620108016, 11835003, 31221003, and 31521063), the Changjiang Scholar Professorship Award (T2015027), the Beijing Brain Initiative of the Beijing Municipal Science & Technology Commission (Z181100001518003), and the Fundamental Research Funds for Central Universities (2019NTST24). We thank the National Center for Protein Sciences at Peking University in Beijing, China, for assistance with MRI data acquisition.

## Author contributions

Tianyuan Lei, Xuhong Liao, Sha Tao, Qi Dong, and Yong He designed the research; Weiwei Men, Yanpei Wang, Mingming Hu, Gai Zhao, Bin Du, Shuping Tan, Jiahong Gao, Shaozheng Qin, Sha Tao, and Qi Dong collected the data; Tianyuan Lei and Xuhong Liao performed the research; Xiaodan Chen, Tengda Zhao, Yuehua Xu, Mingrui Xia, Xiaochen Sun, Yongbin Wei and Qian Wu provided technical assistance; and Tianyuan Lei, Xuhong Liao, Tengda Zhao, Jiaying Zhang and Yong He wrote the paper.

The authors report no biomedical financial interests or potential conflicts of interest.

## Supplementary Information

### Supplementary Methods

#### Participants

We employed a longitudinal rsfMRI dataset consisting of scans taken from 360 typically developing children (F/M = 163/197, 6 to 14 years, 643 scans in total) collected by the Children School Functions and Brain Development project (Beijing Cohort). The children included in this study were cognitively normal, according to a well-validated Chinese standardized cognitive ability test (Dong & Lin, 2011). Participants were excluded if they had a history of neuropsychiatric illness, psychoactive drug use, significant head injuries or significant physical illness, and were not permitted to take drugs, coffee or tea on the day of scanning. Some of these children underwent multiple sessions of multi-modal MRI imaging (T1, T2, rsfMRI, etc.) with an interval of approximately one year between each session. Each rsfMRI scan was assessed against strict quality control criteria before being included in the final dataset. As a result, 152 rsfMRI scans were excluded due to field map error (N = 6), excessive head motion (N = 94), excessive “bad” time points (N = 12) and T1 artifacts (N = 40). Thus, after quality control screening, 491 rsfMRI scans from 305 children (F/M = 143/162, ages 6 to 14 years) were retained for use in our study. These were made up of 3 scans from 47 children (F/M = 31/16), 2 scans from 92 children (F/M = 47/45), and a single scan from 166 children (F/M = 65/101). This dataset was used to explore the longitudinal development of brain network dynamics between childhood to adolescence (Fan et al., 2020). For comparison purposes, we also collected an rsfMRI dataset comprised of scans from 62 healthy young adults (F/M = 37/25, 18 to 29 years), which was acquired using an identical scanner and scanning protocols as those used to obtain the child dataset. After applying the same quality control criteria, data from one participant was excluded due to field map error. The final adult dataset was thus made up of rsfMRI data from 61 adults (F/M = 37/24, ages 18 to 29 years). The study was approved by the Ethics Committee of Beijing Normal University, and written informed consent was obtained from all the participants or their parents/guardians.

#### Image data preprocessing

Resting state fMRI data from child participants was preprocessed using SPM12 (https://www.fil.ion.ucl.ac.uk/spm) and DPABI 3.0 (Yan et al., 2016). Specifically, we first removed the first ten volumes and performed slice-timing correction. Next, a field map correction was applied to remove geometric distortion, which was then followed by a head motion correction. We also estimated the mean framewise displacement (FD) (Power et al., 2012) across time for each scan. Ninety-four scans were excluded due to excessive head motion (i.e., translation > 3 mm, rotation > 3°, or mean FD > 0.5 mm). The motion-corrected functional images were co-registered with individual T1 images and then spatially normalized to a custom template using a unified segmentation algorithm (Ashburner & Friston, 2005) by applying the following procedures: individual T1 images were first segmented into gray matter, white matter and cerebrospinal fluid tissue maps using Chinese Pediatric Atlases (CHN-PD) (6-12 years) (https://www.nitrc.org/projects/chn-pd) (Zhao et al., 2019) as the reference for segmentation. Selecting pediatric atlases specific to Chinese children improves the accuracy in spatial deformation of pediatric brain images. The resulting gray matter, white matter, and cerebrospinal fluid maps were separately averaged across all scans to generate custom tissue templates. T1 image segmentation was then repeated, this time using the custom tissue templates as the reference for segmentation. Subsequently, all individual functional images were spatially normalized to the custom space by applying the transformation parameters estimated during the second T1 segmentation, then resampled to 3-mm isotropic voxels and spatially smoothed using a Gaussian smoothing kernel (full-width at half maximum = 4 mm). Next, we performed linear detrending and nuisance signal regression. During the latter, Friston’s 24 head motion parameters (Friston et al., 1996), “bad” time points with FD above 0.5 mm, and white matter, cerebrospinal fluid and global brain signals were included as covariates. Finally, we used temporal band-pass filtering (0.01-0.1 Hz) to reduce low-frequency drifts and high-frequency physiological noise.

Preprocessing of the adult data was undertaken using the same procedures as that performed on the child functional images, with the exception of the spatial normalization process. The adult functional images were spatially normalized to the Montreal Neurological Institute (MNI) space, with prior white matter, gray matter, and cerebrospinal fluid templates from SPM12 as the normalization reference.

#### Gene data preprocessing

We used brain-wide gene expression data publicly available from the Allen Human Brain Atlas (http://human.brain-map.org/) (Hawrylycz et al., 2012) to conduct our connectome-transcriptome association analysis. This atlas contains tissue samples from six donors, of which samples from two donors cover the whole brain, and samples from the other four donors cover only the left hemisphere. A total of 3,702 tissue samples are included in the atlas, along with their accurate MNI coordinates. Genetic data provided in the atlas has undergone minimal preprocessing in accordance with a white paper published on the Allen Brain Atlas website (http://help.brainmap.org/display/humanbrain/Documentation).

Using this data, we carried out further preprocessing by first removing the samples located in the brain stem and cerebellum, leaving a total of 2,748 samples. Following this, we used the Re-annotation toolkit (Arloth et al., 2015) to re-annotate probe gene names, and removed 10,521 probes with missing Entrez IDs. Next, we used the intensity-based filtering method (Arnatkevičiūtė et al., 2019) to filter the data with reference to background noise intensity. 71% of genes within the same sample have been reported as having been measured with at least two probes (Arnatkevičiūtė et al., 2019). For each of these genes, its expression level within a sample was obtained by averaging the expression values across all detecting probes. After the above procedures were performed, each sample contained expression level data for 15,745 genes. We then undertook normalization of this gene expression data using the scaled robust sigmoid (SRS) algorithm (Fulcher et al., 2013) to minimize the impact of outliers.

Although the Allen Institute has employed strategies to reduce individual differences, further normalization of the gene expression data was still required. We first performed cross-gene normalization within each sample, followed by cross-sample normalization for each gene within the same donor, in order to correct for measurement errors and discrepancies between different samples and donors (Burt et al., 2018). Next, we matched the MNI coordinates of each sample to the random-1024 parcellation scheme of the adult group using the nearest-point search algorithm (Barber et al., 1996), assigning each sample to one of the brain nodes. For each node, the expression data of each gene was obtained by first averaging the data across the samples within the same donor and then averaging the nodal expression data across donors. This dual-averaging operation reduces the effect of inhomogeneous spatial distributions in samples across different donors and ensures that each donor makes the same contribution to the gene expression profile. The gene preprocessing described above was performed by referencing the code at https://github.com/BMHLab/AHBAprocessing (Arnatkevičiūtė et al., 2019).

## Supplementary figures

**Figure S1.**
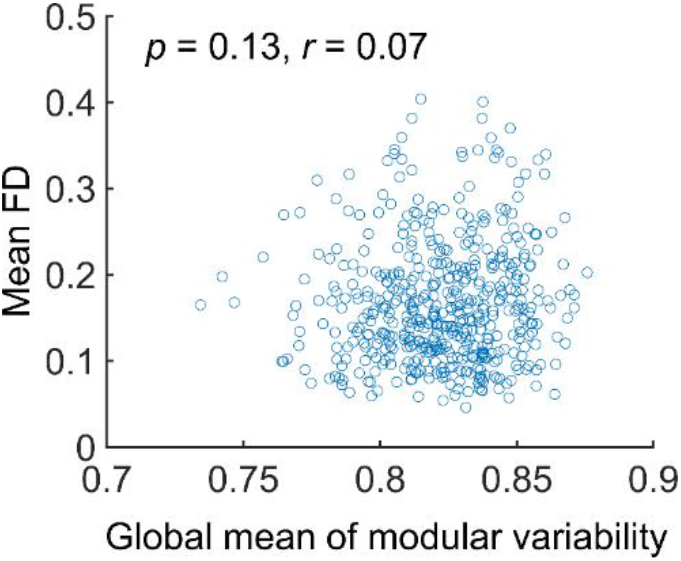
Relationship between brain network dynamics and head motion in children. Based on a Pearson’s correlation analysis across rsfMRI scans, there was no significant correlation between the mean modular variability of the whole brain and the mean FD across time windows (*p* > 0.05). Every circle represents one child rsfMRI scan. FD, framewise displacement.

**Figure S2.**
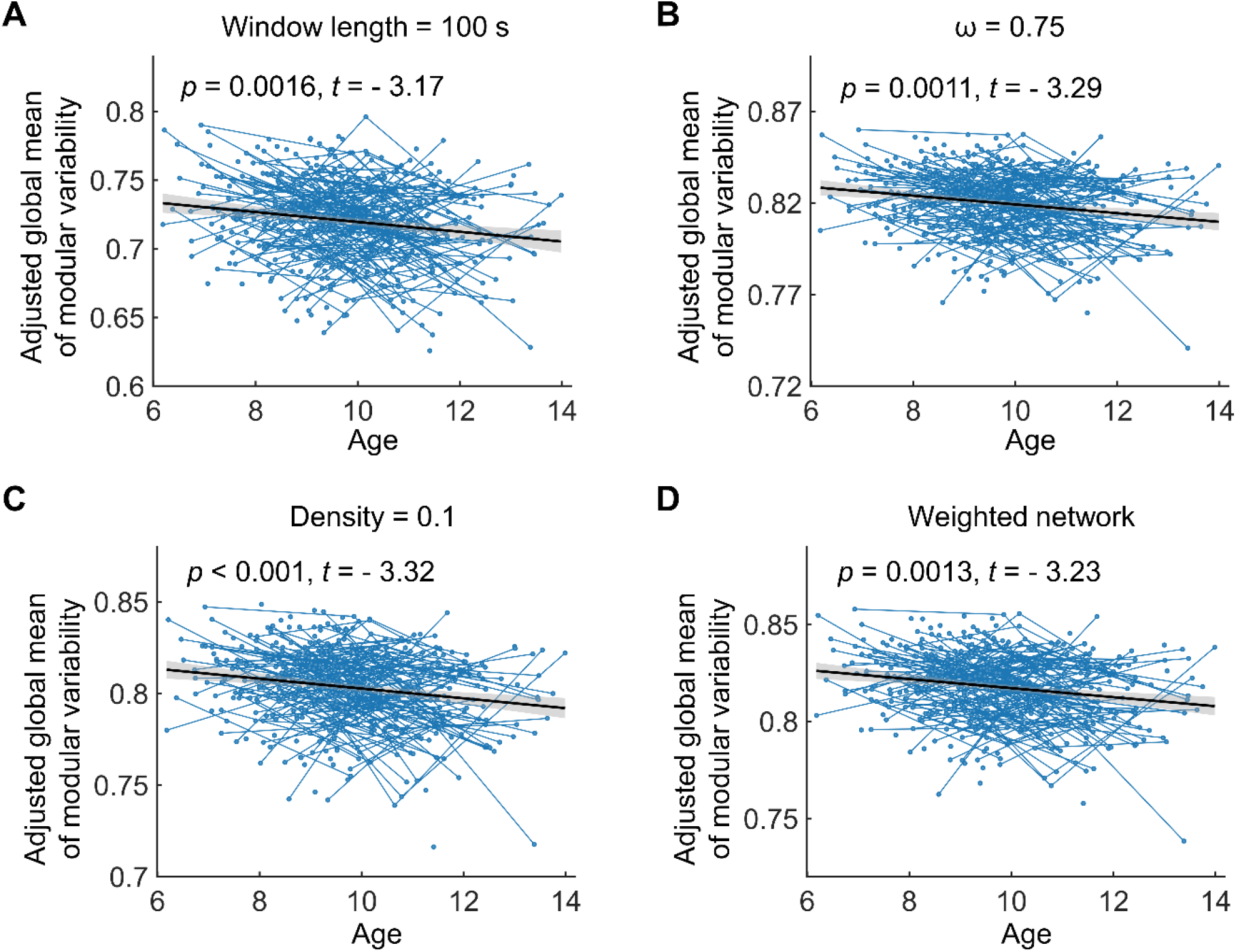
Age-dependent changes in global brain network dynamics of children under different network analysis strategies. **(A)** Sliding window length = 100 s. **(B)** Temporal coupling parameter *ω* = 0.75. **(C)** Binary networks with a network density of 10%. **(D)** Weighted networks with a network density of 5%. In each alternative analysis, all network analysis parameters were set to be the same as those in the main analysis, except for the parameter of interest. Age effects were estimated using a mixed effect model. The adjusted value denotes the measure of interest corrected for sex, head motion, and random age effects. In all cases, the mean modular variability of the brain significantly decreased with age between childhood to adolescence.

**Figure S3.**
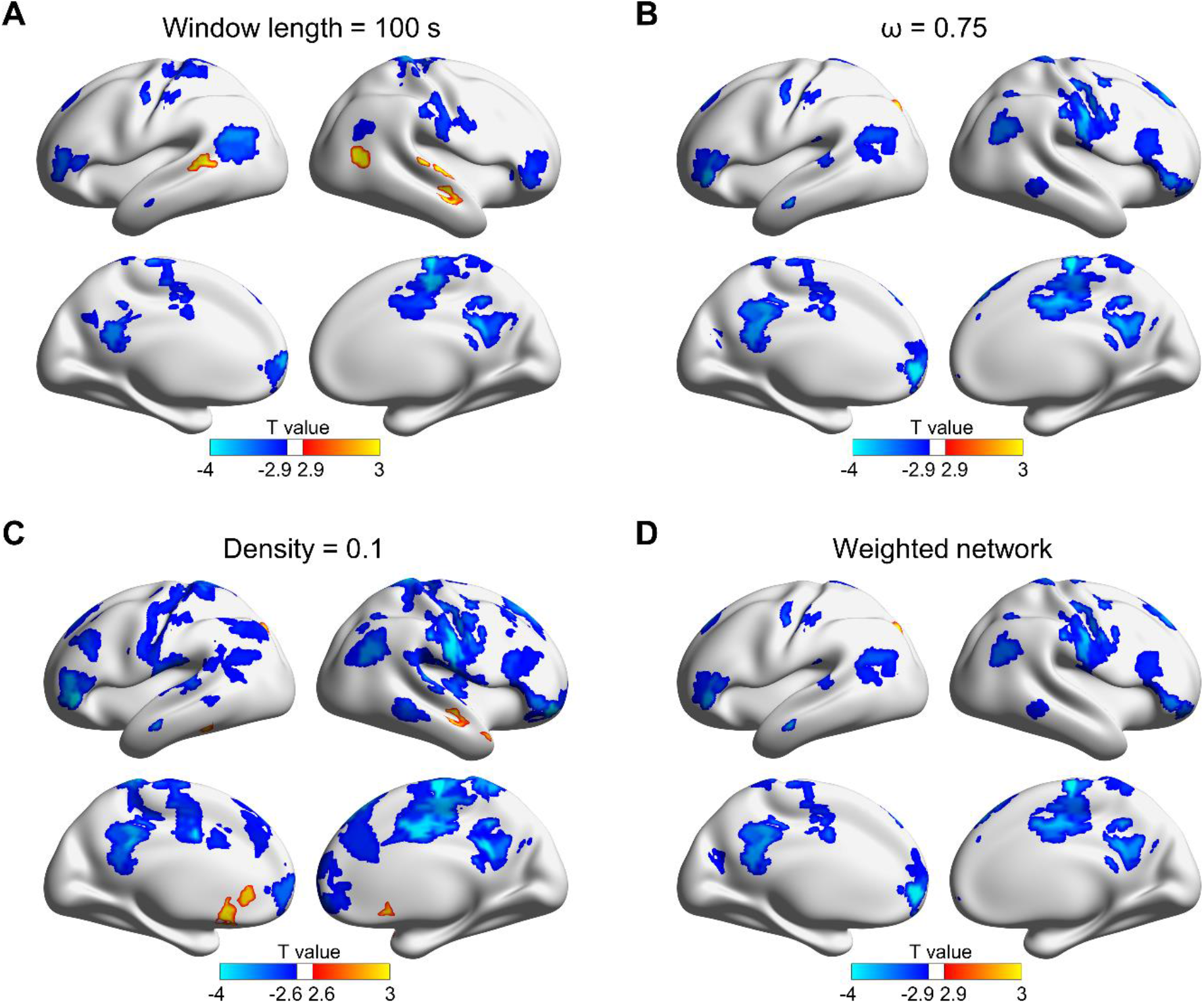
Developmental changes in regional brain network dynamics of children under different network analysis strategies. **(A)** Sliding window length = 100 s. There were 64 brain nodes showing significant age-related changes in regional network dynamics (FDR-corrected *p* < 0.05), of which 59 regions showed a linear decrease with age. **(B)** Temporal coupling parameter *ω* = 0.75. There were 74 brain regions showing significant age-related changes in regional network dynamics (FDR-corrected *p* < 0.05), of which 73 regions showed a linear decrease with age. **(C)** Binary networks with a network density of 10%. There were 178 brain regions showing significant age-related changes in regional network dynamics (FDR-corrected *p* < 0.05), of which 171 regions showed a linear decrease with age. **(D)** Weighted networks with a network density of 5%. There were 75 brain regions showing significant age-related changes in regional network dynamics (FDR-corrected *p* < 0.05), of which 74 regions showed a linear decrease with age. Age effects were estimated using a mixed effect model. Regional network dynamics was measured by the modular variability of network nodes. Note that significant regions in these four cases showed high levels of spatial overlap with those observed in the main results.

**Figure S4.**
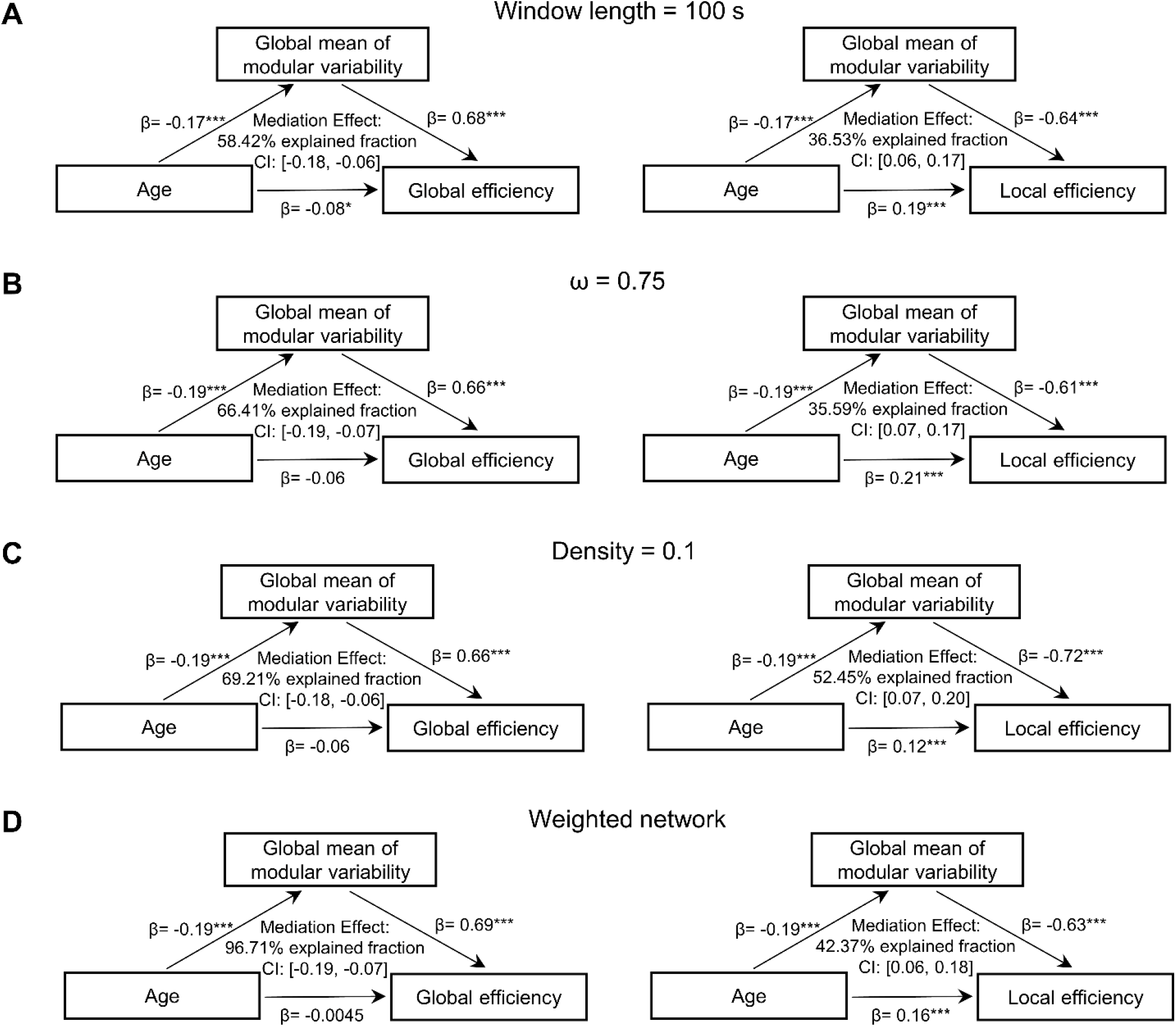
Mediation effect of brain network dynamics on the development of global efficiency and local efficiency in children under different network analysis strategies. **(A)** Sliding window length = 100 s. **(B)** Temporal coupling parameter *ω* = 0.75. **(C)** Binary networks with a network density of 10%. **(D)** Weighted networks with a network density of 5%. In general, the age-related reduction in functional module dynamics significantly mediated the age-related decrease in global efficiency and the age-related increase in local efficiency of brain networks, regardless of the network analysis strategy applied.

**Figure S5.**
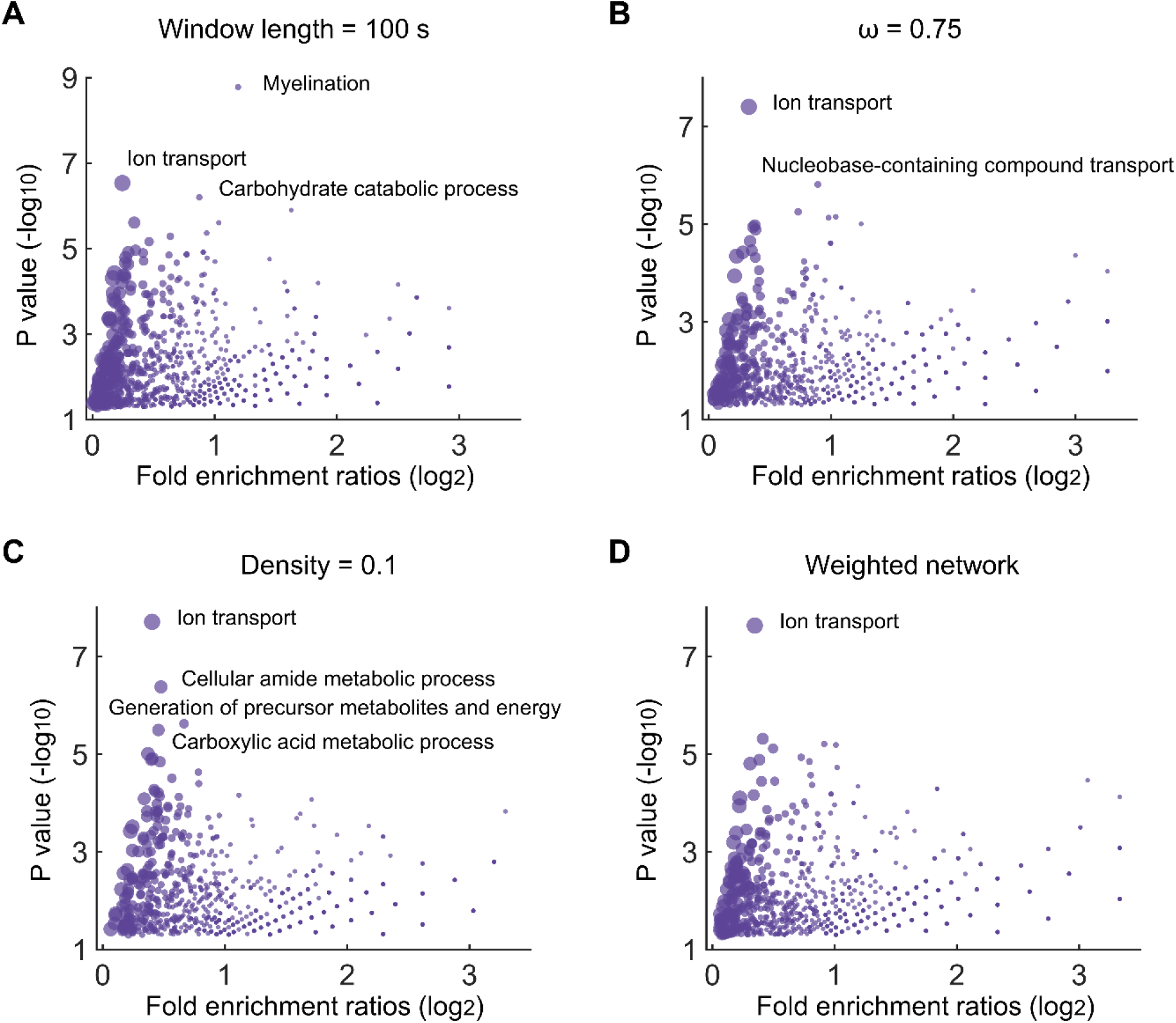
Gene ontology (GO) terms of biological processes associated with genes showing significant positive correlations with developmental changes in brain network dynamics under different network analysis strategies. **(A)** Sliding window length = 100 s. **(B)** Temporal coupling parameter *ω* = 0.75. **(C)** Binary networks with a network density of 10%. **(D)** Weighted networks with a network density of 5%. Gene ontology annotation analyses were conducted using the ToppGene Suite (https://toppgene.cchmc.org/). Dots marked with text represent significant GO terms obtained with correction applied for multiple comparisons (FDR-corrected *p* < 0.05), and the remainder represent GO terms obtained where no correction was applied (uncorrected *p* < 0.05). The dot size represents the number of genes that overlap with the corresponding GO term. In **(A)**, in addition to the biological processes annotated in the plot, there were 6 significant GO terms unmarked due to space limitations. These are: i) cortical actin cytoskeleton organization; ii) regulation of catabolic process; iii) glial cell development; iv) ribonucleoside diphosphate metabolic process; v) cellular carbohydrate metabolic process; and vi) generation of precursor metabolites and energy. Notably, the GO term ion transport was consistently observed in the enrichment analysis under different network analysis strategies.

## Supplementary Tables

**Table S1.**
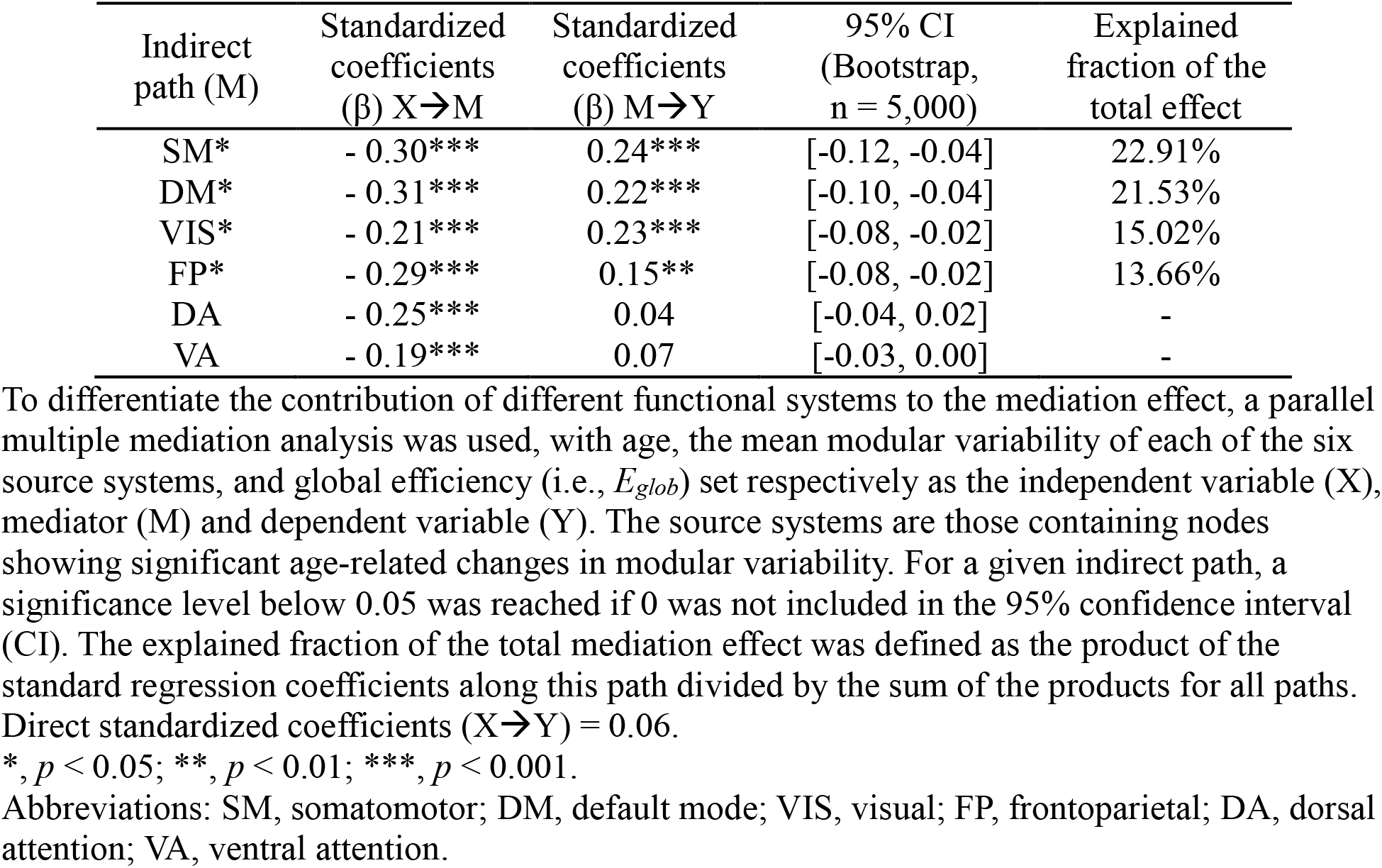
Estimated parameters in the parallel multiple mediation analysis for global efficiency between childhood to adolescence

| Indirect path (M) | Standardized coefficients ($\beta$) $X \rightarrow M$ | Standardized coefficients ($\beta$) $M \rightarrow Y$ | 95% CI (Bootstrap, $n = 5,000$) | Explained fraction of the total effect |
| --- | --- | --- | --- | --- |
| SM* | - 0.30*** | 0.24*** | [-0.12, -0.04] | 22.91% |
| DM* | - 0.31*** | 0.22*** | [-0.10, -0.04] | 21.53% |
| VIS* | - 0.21*** | 0.23*** | [-0.08, -0.02] | 15.02% |
| FP* | - 0.29*** | 0.15** | [-0.08, -0.02] | 13.66% |
| DA | - 0.25*** | 0.04 | [-0.04, 0.02] | - |
| VA | - 0.19*** | 0.07 | [-0.03, 0.00] | - |

To differentiate the contribution of different functional systems to the mediation effect, a parallel multiple mediation analysis was used, with age, the mean modular variability of each of the six source systems, and global efficiency (i.e., $E_{glob}$ ) set respectively as the independent variable (X), mediator (M) and dependent variable (Y). The source systems are those containing nodes showing significant age-related changes in modular variability. For a given indirect path, a significance level below 0.05 was reached if 0 was not included in the 95% confidence interval (CI). The explained fraction of the total mediation effect was defined as the product of the standard regression coefficients along this path divided by the sum of the products for all paths. Direct standardized coefficients ( $X \rightarrow Y$ ) = 0.06.
\*, $p < 0.05$ ; \*\*, $p < 0.01$ ; \*\*\*, $p < 0.001$ .
Abbreviations: SM, somatomotor; DM, default mode; VIS, visual; FP, frontoparietal; DA, dorsal attention; VA, ventral attention.

**Table S2.**
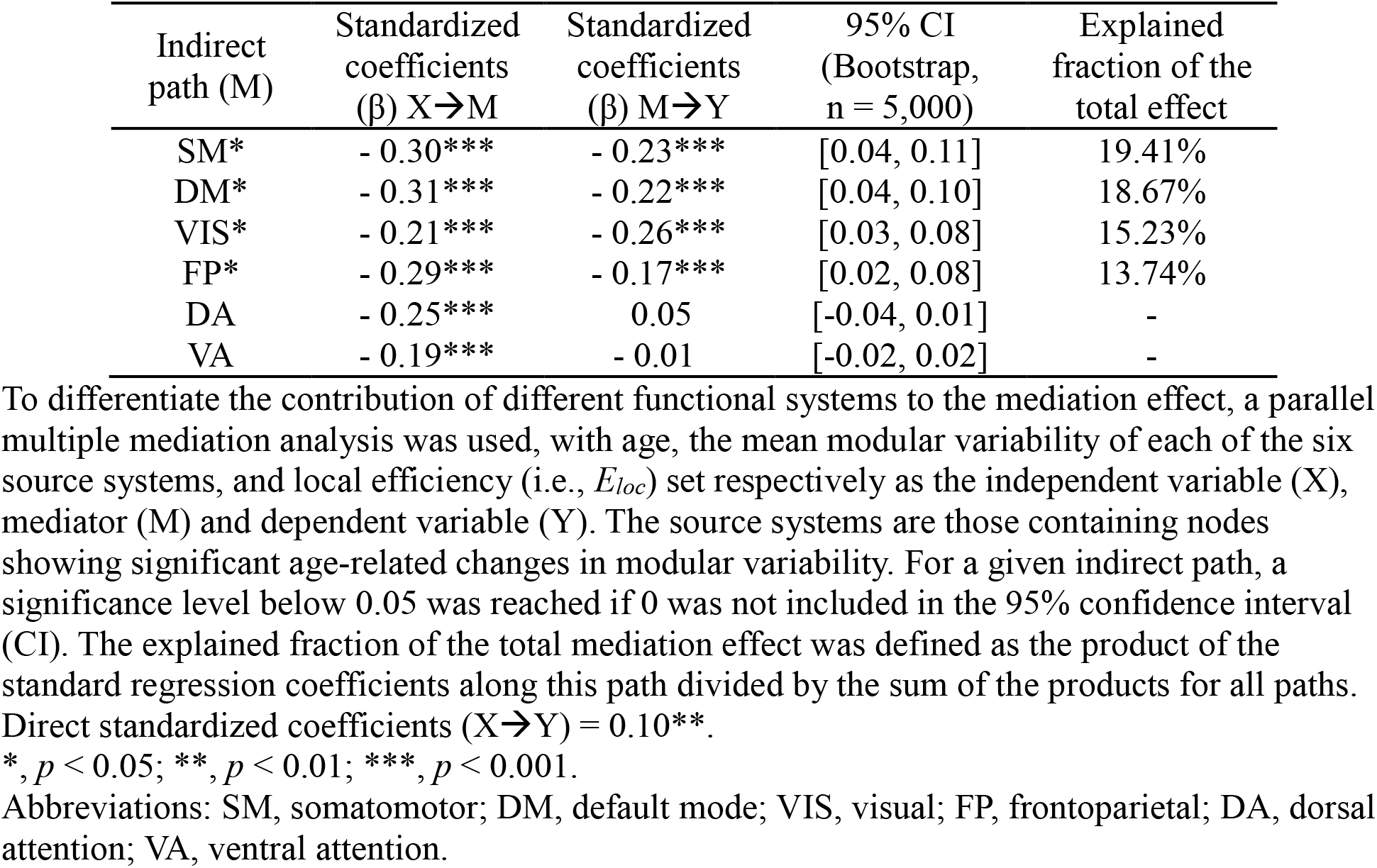
Estimated parameters in the parallel multiple mediation analysis for local efficiency between childhood to adolescence

**Table S3**. List of genes showing a significant positive correlation with the developmental change in modular variability in the main analysis (Table S3. xlsx). This table is available at https://github.com/helab207/Development-of-brain-module-dynamics.

**Table S4**. Enrichment analysis results (i.e., GO terms of biological process) for genes with a significant negative correlation with the developmental changes in modular variability in the main analysis (Table S4. xlsx). This table is available at https://github.com/helab207/Development-of-brain-module-dynamics.

